# The role of many-to-one mapping of vertebral form to function in Psittaciform tripedal locomotion

**DOI:** 10.1101/2024.02.29.582755

**Authors:** Anna R. Stuart, Michael C. Granatosky, Ryan N. Felice, Ryan D. Marek

## Abstract

Parrots highlight the functional diversity of the avian neck by contributing to a range of behaviors, including arboreal locomotion. The parrot neck is used alongside the beak and hindlimb to allow them to successfully navigate arboreal habitats via tripedal locomotion. Whether specific morphological characteristics of the neck enable this behavior are currently unknown. By combining geometric morphometrics with phylogenetic comparative methods we investigate the factors correlate with shape variation in the cervical vertebrae of parrots. We find that phylogeny, allometry, integration, diet and tripedal locomotion all have a significant influence on the morphology of psittaciform cervical vertebrae. However, the influence of diet and tripedal locomotion is weak, with a high degree of morphospace overlap existing between dietary and neck use groups. Additionally, we find no evidence of convergence in parrot neck morphology due to the incidence of tripedal locomotion or dietary specialization. We thus conclude that changes to the neuromuscular control of the neck, not morphological adaptations, are primarily responsible for tripedal locomotion in parrots. We argue that many-to-one mapping of form to function allows parrots with similar neck morphologies to participate in a range of behaviors, and this may be a common feature amongst all birds.

## Introduction

The avian forelimb is characterised by its highly specialised anatomy that allows for powered flight. As a consequence of this derived structure and function, it is generally maladapted for many functions outside of this key innovation (1–3). As such a large number of functions that were ancestrally performed by the forelimb are instead shifted to the craniocervical system (i.e., beak, head, and neck) in birds. Thus, the neck of birds functions across a broad range of activities as a ‘surrogate forelimb’ (3–5). Psittaciformes (parrots) exemplify the functional diversity of the avian cervical column as the necks of this group participate in feeding, preening, tool use and most spectacularly, locomotion (6–8). By utilising the neck and head as a third ‘propulsive limb’ parrots are able to ascend vertical and traverse horizontal substrates by co-opting the craniocervical system to function within a cyclical tripedal gait pattern (6,8,9). This adaptation of the craniocervical system to function as a propulsive limb appears to be an evolutionary novelty unique to parrots, and investigating this phenomenon may offer insight into how the head and neck can be exapted to actively participate in locomotion.

The use of non-appendicular appendages in locomotion is well-documented across tetrapods. If one of these appendages displays an interaction between the animal’s mass and the substrate then it can be defined as ‘effective limb’ (10). There are numerous examples of portions of the axial column (particularly the tail) acting as an effective limb, however many of these are interacting with the substrate in an incidental manner (e.g. tail dragging (11–13)), or acting as a stabiliser/brace (6). Propulsive limbs (i.e., effective limbs that are used for propulsion (6)) are much rarer in tetrapods and are often limited to the involvement of a tail in powered pentapedal gaits (14,15). Recent work has highlighted the extraordinary ability of Psittaciformes to use the craniocervical system as a propulsive limb, with the beak, neck and hindlimbs generating more relative tangential substrate reaction forces than the forelimbs of humans and primates during vertical climbing (6,8,16). A large proportion of this propulsive force must be driven by the neck as the thoracolumbar spine of birds is adapted for stiffness and stability, not force production (6). This has led to the hypothesis that either the neuronal (6,16) or musculoskeletal system of the psittaciform neck has undergone radical changes to accommodate tripedal locomotion. If the adaptation is neuronal then many-to-one mapping of form to function may be a key component of neck evolution in parrots, allowing novel forms of locomotion to evolve rapidly without the need for large shifts in neck morphology across the entire cervical spine. Parrots may have utilised many-to-one mapping over the course of their evolutionary history to allow them to use their neck in a locomotory capacity without sacrificing the diverse functionality that the ‘surrogate forelimb’ provides (4). If the adaptation is musculoskeletal, then these changes can be observed by quantifying relationships between vertebral morphology and neck usage in parrots.

Many-to-one mapping is a vital concept within evolutionary biology as it can result in a decoupling of functional diversity from morphological diversity within a clade (17,18). This decoupling allows for one morphological trait to accommodate multiple functions, and highlights that the morphological form does not always directly correlate with function (17,19). Many-to-one mapping may be exemplified by the avian neck as it performs a wide range of functions despite a similarity in its overall structure between species, and this is evidenced by recent work which observed that the regional modularity of the avian neck is conserved across many species, only adapting to highly specialised functions such as carnivory (3,20). Here we investigate whether the psittaciform neck breaks this pattern of conservatism in avian neck morphology in order to adapt to function during locomotion, or if many-to-one mapping and changes to neuronal control of parrot neck musculature are responsible for this novel behaviour.

Parrots have long been known to engage in climbing behaviour and tripedal locomotion during climbing is thought to be a universal feature of Psittaciformes (21–23). Yet this ubiquity of tripedal locomotion in parrots is anecdotal (21) and the extent of beak-assisted climbing in this clade has never been quantified. By using a framework set out by recent research into avian foot use (24) we can leverage enormous photographic databases of parrots identified to species level in order to quantitatively determine the incidence of beak-assisted climbing across a broad range of extant Psittaciformes. We then use the incidence of climbing data to assess any potential form-function relationships between neck vertebral morphology and beak-assisted climbing. As the psittaciform neck provides much of the propulsive force associated with climbing (6) we hypothesize that a significant portion of vertebral shape variation will be governed by the incidence of beak-assisted climbing. We may further speculate that the selective pressures potentially imposed by beak-assisted climbing upon the neck leads to morphological convergence of the cervical spine in groups that are frequently observed carrying out this behaviour. Prior work can be used as a basis for this hypothesis, and has observed that non-psittaciform birds that share specialised neck kinematics (e.g. carnivorous birds) display similar patterns of gross neck morphology (3). Phenotypic Integration, the co-evolution of anatomical traits, is an important facilitator of morphological diversity and has been found to be an important component of avian skull evolution (25,26). Integration often occurs when multiple anatomical systems function together during a particular behaviour (27–29), and recent data suggests it is a commonplace amongst extant avians (30). As tripedal locomotion involves the neck and hindlimb working in tandem, we also hypothesize that neck-hindlimb integration will significantly influence morphological variation of neck vertebrae across Psittaciformes.

Here we use a combination of geometric morphometrics, phylogenetic comparative methods and behaviour quantification to investigate the relationship between neck usage (with a focus on beak-assisted climbing) and neck vertebral morphology across a phylogenetically broad selection of extant parrots. We also assess the relative contributions of other factors that have previously been found to significantly influence the morphological variability of neck vertebral morphology, such as body size, phylogeny, ecology, and morphological integration. Finally, we will use recently established convergence metrics to test for convergence in the vertebral morphology parrots that frequently undertake beak-assisted climbing.

## Methods

### Specimen details

We analysed the morphology of the cervical vertebrae, head, forelimb and hindlimb of 48 species of extant parrots (Supplementary Table 1). Neck use behavior was also studied for 44 of 48 of these species. Six of the 48 species were scanned at UCL with a Nikon XT H 225 microCT scanner. Scan data for the remaining 42 species were downloaded from MorphoSource (see Supplementary Table 1 for MorphoSource ID numbers). Dietary and foraging guilds for these species were taken from the AVONET database (31). Phylogenetic trees were acquired from www.birdtree.org and pruned to include only the 48 species included in this study.

### Digitisation and 3D geometric morphometrics

Scans for all 48 species were segmented in Amira 3D (version 2021.1, Visualization Science Group, Thermo Fisher Scientific) and the digital models outputted were cleaned and further processed in MeshLab (32). To account for the variation in total cervical vertebral counts between species we analysed 2 homologous vertebrae (the second cervical vertebrae, C2, and the last cervical vertebrae) as well as 3 ‘functionally homologous’ vertebrae (vertebrae at 25%, 50% and 75% along the cervical column) (20,33,34). A vertebral landmark scheme consisting of 22 fixed landmarks (Supplementary Figure 1, Supplementary Table 2) was constructed based on schemes from prior avian vertebral morphometric studies (3,20,35,36). We next calculated head volumes by subjecting digital skull models for each species to an α-shape fitting algorithm that is part of an in-house modified version of the ‘alphavol’ package for MatLab (3,37). These volumes were then multiplied by the weighted mean densities of soft tissues within the skull (approximated to the density of water, 997 kg/m^3^). This method may overestimate head mass as it does not model the degree of pneumatic bone and soft tissue within each skull, but such granularity was outside the scope of this study. Head mass was preferred over head shape as head shape often poorly correlates with ecology in birds (25,26,38). We then measured limb element lengths digitally in Geomagic Wrap (Geomagic, United States). Forelimb elements measured include the coracoid, scapula, humerus, radius, ulna and carpometacarpus. Hindlimb elements measured include the femur, tibiotarsus and tarsometatarsus. We size-corrected limb measurements using the following formula: limb element length/body mass^0.33^. We also size-corrected head mass by calculating the percentage of total body mass. Scaling equations based on the humeral articulation facet of the coracoid were used to estimate body masses (39).

### Assessment of neck use behavior

To assess the extent to which the neck was being used by parrots, we devised a scoring system that noted the presence or absence of four neck functions assumed to put relatively high loads on the neck [i.e., beak assisted climbing (6,9,16), forceful flexion and extension (40–42), object carrying)]. We used the Macaulay Library (https://www.macaulaylibrary.org) to search for images of parrots performing these neck use behaviors following protocols from prior literature (24). Up to 2000 images were searched for each of the 48 species in the study (24) (see Supplementary Table 3 for behavior scorings and incidence data). We assessed the relationship between number of photos searched and neck use score for each behavior and found it to be significant for each of the four neck behaviors, as well as for the overall presence or absence of absolute neck use. To accommodate for this we use an incidence of behavior metric that accounts for the number of photos searched. Initially we observed a significant relationship between this incidence metric and the number of photos searched, but after the removal of 4 outliers (species with an unusually high incidence of beak assisted climbing despite < 50 photos searched per species) there was no significant correlation (p < 0.05).

It should be noted that our method for scoring neck function was a necessity owing to the lack of available quantitative information on neck use in parrots (43). Generally, this approach is synonymous with scan sampling, which provides an unbiased assessment of the activity budget of an animal, but suffers from missing rare or uncommon behaviors (44,45). Indeed, this is confirmed with the relatively low occurrence of neck use observed in parrots in this study (see below). For rare behaviours, focal animal or specific behaviour sampling would be most appropriate (44,45). However, no studies have yet to quantify the positional behaviour of parrots, let alone any non-mammalian species, utilizing such methodology.

### Statistical analysis

#### Morphological analysis

We first subjected landmark data to Procrustes superimposition. We then performed morphological analyses on a pooled dataset that contained all vertebrae (C2, C25%, C50%, C75% and the last cervical vertebrae) for all species, as well as separate analyses for each vertebral level. This allowed us to study morphological evolution across the entire parrot neck and at the level of individual vertebrae across the cervical spine (20). We calculated multivariate Blomberg’s K (K_mult_) using the function physig() within the ‘geomorph’ R package to estimate the impact of phylogeny on the morphological variation of parrot cervical vertebrae (46). To assess the relationship between vertebral shape variation and ecological parameters, neck use behaviors and body mass we utilized phylogenetic multivariate ANOVAs (pMANOVAs). A reduced dataset was used in the neck use behavior pMANOVAs as we identified some outlier species (see prior methods section, ‘Assessment of neck use behavior’). We determined significant differences in morphology between ecological groups using post-hoc pairwise tests with the ‘pairwise’ function in the ‘RRPP’ package in R.

We identified the degree of head-neck, neck-forelimb and neck-hindlimb integration using a phylogenetic two-block partial least-squares (2BPLS) analysis in the R package ‘geomorph’ (46). We then used the ‘compare.pls’ function (as part of the ‘geomorph’ R package) to search for significant differences in effect sizes across integration tests. Next, we combined all measurements of individual forelimb and hindlimb elements into a single forelimb or hindlimb matrix prior to inclusion in any 2BPLS analyses. Following this, we carried out pMANOVAs and 2BPLS tests for a pooled vertebral dataset and for each individual vertebral level. In order to assess differences in morphological shape change across the entire neck we applied Phenotypic Trajectory Analysis (PTA) (47). PTA plots a trajectory through shape space for a particular group (in this case for a particular dietary or foraging category) by connecting the mean shape of a particular vertebrae in a sequential chain from C2 to the last cervical vertebrae. This allows for the statistical assessment of differences between certain groups in the pattern of sequential vertebral shape change across the entire neck.

We produced phylomorphospace plots of PC1 and PC2 for each individual cervical vertebrae to visualize differences in morphospace occupation between ecological groups and between disparate incidences of neck use behavior values. We focused on exploring these axes because for each vertebra, the subsequent PC axes each explained < 30% of the overall variation in each ordination. We used these phylomorphospace plots to inform our hypotheses for morphological convergence analyses. These visualizations allowed us to identify focal taxa that share the same ecology or neck use traits and also occupied a distinct area of morphospace and were thus candidates for convergent evolution. We then tested for morphological convergence using the Ct1-Ct4 metrics using the ‘calcConvCt’ and ‘calcSigCt’ functions in the R package ‘convevol’ (48,49).

#### Behavior analysis

Following the protocol of previous work (24) we first tested for any correlation between the number of photos studied versus the overall occurrence of neck use behavior with a PGLS using the ‘gls’ function of the R package ‘nlme’. The absolute occurrence of neck use displayed a significant correlation with the number of media analysed, and as such we accounted for the number of media by using an incidence of neck use metric. We used Phylogenetic ANOVAs to assess potential correlations between the incidence of neck use behaviors and ecological categories, body mass and head mass. We visualized the distribution of the incidence of neck use behavior across our sample by mapping this trait onto a phylogenetic tree using the ‘contMap’ function in ‘phytools’ (50).

## Results

### Morphological variation of parrot neck vertebrae

Three out of five of the studied cervical vertebrae occupy distinct areas of morphospace when all vertebrae are projected onto the same morphospace, with only C50% and C75% displaying any level of overlap (Supplementary Figure 4). Principal component (PC) 1 accounts for 38.15% of the morphological variation and higher PC1 scores are associated with a lengthening of the centrum and vertebral arch, a decrease in size of the neural spine, a widening of the neural canal and vertebral arch, and an increase in length of the costal processes (Supplementary Figure 4). PC2 accounts for 27.63% of the morphological variation, with higher PC2 scores corresponding to a decrease in centrum and vertebral arch length, an increase in neural spine height, a narrower neural canal, more robust prezygopophyses and transverse processes, a widening of pre- and post-articular facets and in increase in the size of the ventral spine (Supplementary Figure 4).

We then ordinated morphological variation for each vertebrae individually, revealing two distinct patterns for either proximal or distal vertebrae. Proximal vertebrae (C50%, C75% and the last cervical vertebrae) tended to display elongated and deeper centrums along the main axis of variation (PC1), whilst centrum length shortened across PC1 for more distal vertebrae (C2 and C25%). PC1 corresponds to between 21.4% (C25%) and 43.78% (C75%) of the vertebral morphological variation, and PC2 corresponds to between 11.27% (C2) and 16.28% (C50%) of vertebral morphological variation (Figure 1). Increases to values of PC1 of C2 vertebrate correspond to a shortening of the centrum and vertebral arch, an increase in height and width of the neural spine, more robust zygapophyses and a reduction in size of both articular facets (Figure 1). An increase in PC2 scores of C2 vertebrae is associated with a shortening and narrowing of both the centrum and vertebral arch alongside an increase in neural spine height (Figure 1). Increases to PC1 scores of C25% vertebrae correspond to a widening and shortening of the centrum and vertebral arch, a small increase in height of both the neural and ventral spines, and a reduction in the height of the postarticular facet (Figure 1). Increases to PC2 values correspond to subtle changes in C25% morphology such as a less posteriorly pronounced ventral lip of the postarticular facet and a slight reduction in costal process length (Figure 1). For middle (C50%) vertebrae, increases in PC1 scores are associated with a slight increase in the length and depth of the centrum, a posterior shift in the position of the neural spine, a narrowing of the transverse processes, shorter postzygapophyses and elongated costal processes (Figure 1). Increases to PC2 scores in middle vertebrae are associated with an increased neural spine height and enlarged transverse processes (Figure 1). Changes in vertebral shape across PC1 in C75% vertebrae correspond to an increased centrum depth, a slight increase in neural spine height, a minor elongation of the costal processes, and a more concave postarticular surface (Figure 1). Increases in PC2 scores in C75% vertebrae are associated with increases to the width of both the transverse processes and the postzygapophyses (Figure 1). For the last cervical vertebrae changes associated with increased PC1 scores correspond to an increase in centrum length and depth, increases to neural and ventral spine height, narrower transverse processes, more dorsally positioned prezygapophyses and narrower postzygapophyses (Figure 1). Shape changes associated with an increase in PC2 scores of the last cervical vertebrae correspond to a more anteriorly positioned postzygapophysis and an enlarged postarticular facet (Figure 1).

**Figure 1.**
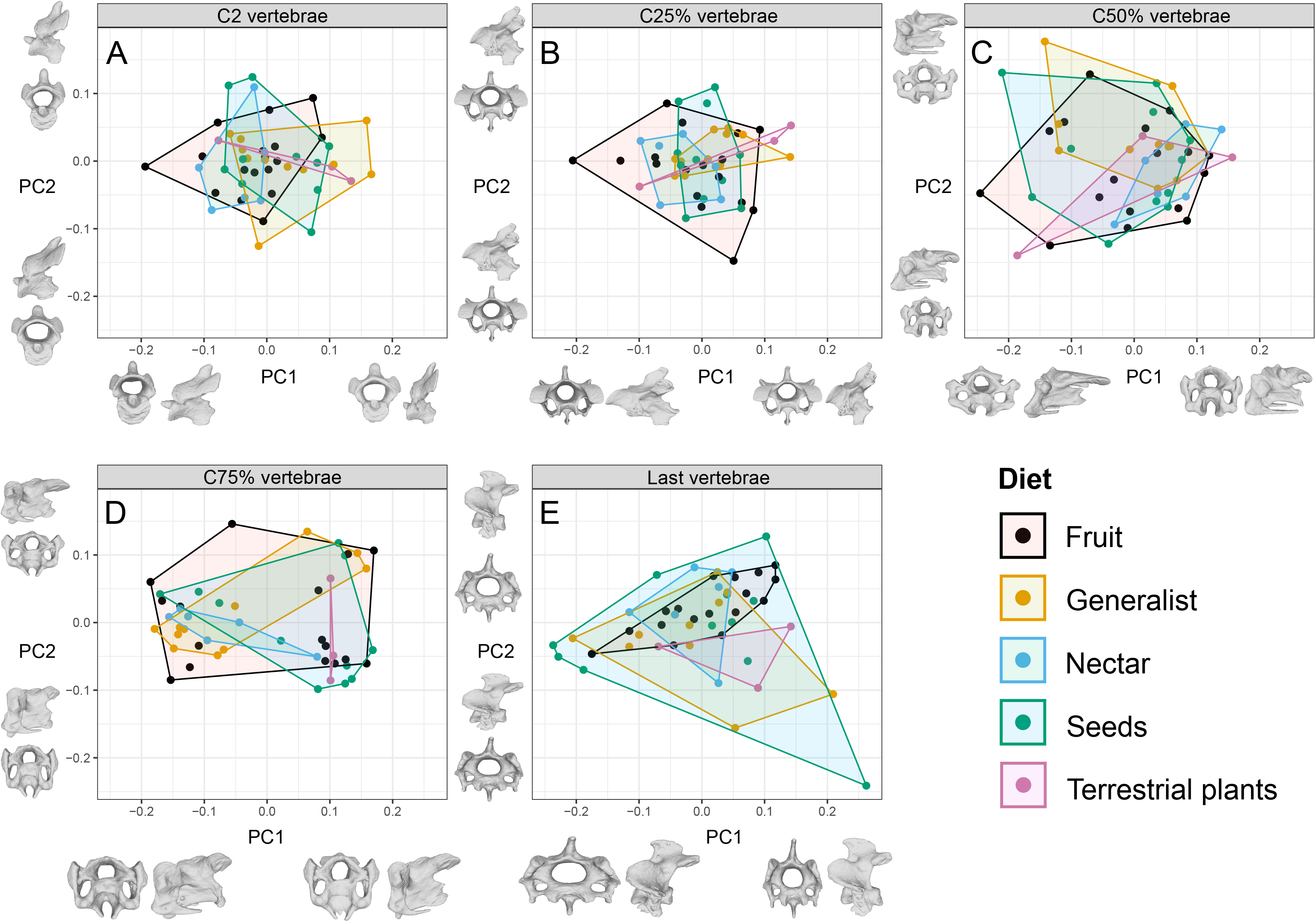
Morphospaces for each of the psittaciform cervical regions studied, grouped by dietary preference. A) C2 cervical vertebrae, B) C25% cervical vertebrae, C) C50% vertebrae, D) C75% vertebrae, E) Last cervical vertebrae. Warped meshes of cervical vertebrae display changes across PC1 and PC2 in anterior (left along PC1, bottom along PC2) view and left lateral (right along PC1, top along PC2) view.

### Allometric and phylogenetic signal in parrot neck morphology

There is a significant relationship between body mass and vertebral shape in the pooled dataset (p = 0.004) as well as in C2 (p = 0.001), C25% (p = 0.002) and the last cervical vertebrae (p = 0.035) (Supplementary Table 4). However, the proportion of morphological variation that can be explained by body mass alone is small and varies between 0.044 (last cervical vertebrae) and 0.075 (C2) (Supplementary Table 4). We calculated K_mult_ to determine the strength and significance of the phylogenetic signal on the morphological variation of parrot cervical vertebrae. A significant (P < 0.05) and weak-to-moderate (lowest K_mult_ C2 = 0.496, highest K_mult_ last = 0.672) phylogenetic signal can be observed in the morphological variation of the pooled vertebral dataset, as well as for all but one (C75%, p = 0.111, K_mult_ = 0.479) of the individual cervical regions.

### Ecological signal in parrot neck morphology

Results from the pMANOVAs suggest significant differences exist in morphology between dietary groups within the pooled dataset (p = 0.004), as well as within C2 (p = 0.004), C25% (p = 0.014) and C75% (p = 0.014) vertebrae (Supplementary Table 4). Diet displayed a weak correlation with vertebral shape across all vertebrae studied (R^2^ between 0.121 in C25% and 0.151 in C75%), yet displayed a comparatively higher coefficient of correlation than body mass (R^2^ 0.045 – 0.075) (Supplementary Table 4). Foraging guild displayed no significant correlation (P > 0.05) with vertebral morphology across any of the vertebral regions studied (Supplementary Table 4). This weak ecological and phylogenetic signal in parrot cervical morphology is reflected in the lack of distinction between ecological and family-level groups in morphospace across all vertebral regions studied.

A post-hoc pairwise test revealed that between dietary groups, terrestrial herbivores and granivores often displayed significantly different vertebral morphologies compared to most other dietary groups (Supplementary Table 5). Terrestrial herbivores often displayed significant differences in vertebral morphology compared to other dietary groups (8 comparisons across C2, C25% and C75%) (Supplementary Table 5). Vertebral morphology was significantly different between herbivores and frugivores in C2 (p = 0.043) and C25% (p = 0.049), between herbivores and generalists in C2 (p = 0.04), C25% (p = 0.04) and C75% (p = 0.017) and between herbivores and nectarivores in C2 (p – 0.012), C25% (p = 0.015) and C75% (p = 0.02) (Supplementary Table 5). Multiple dietary groups also displayed significant differences in vertebral morphology when compared to granivores, including nectarivores (C2 p = 0.026 and C25% p = 0.044) and generalists (C75%, p = 0.032) (Supplementary Table 5). Terrestrial herbivores had predominantly taller neural spines than other dietary groups across C2, C25% and C75%, a shortened centrum in C25% and a deeper centrum in C75%. Granivores also display a taller neural spine across C2, C25% and C75%, as well as comparatively deeper centra in C2 and C75%.

PTA detected that across all dietary, foraging and neck-use groupings, patterns of whole-neck morphological variation were only significantly different between a select few dietary groups (Supplementary Table 6). We tested for pairwise differences between dietary, foraging and phylogenetic groupings separately and found only 4 dietary pairwise comparisons to be significant (Supplementary Table 6, Supplementary Figure 4). Trajectory shape was significantly different between terrestrial herbivores and generalists (p = 0.02), as well as between terrestrial herbivores and granivores (p = 0.035) (Supplementary Table 6). The magnitude of differences between trajectories was significant between generalists and frugivores (p = 0.035), as well as between generalists and granivores (p = 0.005) (Supplementary Table 6).

### The influence of neck use on parrot neck morphology

Our survey of neck use across 44 species of parrots (excluding outliers, see Methods) revealed that across 40,893 images studied, the incidence of absolute neck use was 0.638%. Across each individual neck use behavior, average incidence varied between 0.450% for beak-assisted climbing and 0.032% for object carrying (Supplementary Table 3). Numerous species (out of 48 total) were never observed performing each behavior and this value ranged between 13 and 14 in absolute incidence of neck use and beak assisted climbing and 40 in forceful extension (Supplementary Table 3). Before evaluating the relationship between the incidence of neck use behaviors and cervical morphology, we assessed the relationship between behavior, body mass, head mass and ecology (diet and foraging guild). Across all behaviors only the overall incidence of absolute neck use was significantly correlated with body mass (p < 0.001) and head mass (p = 0.002), beak assisted climbing was also significantly correlated with head mass only (p = 0.016) (Supplementary Table 7).

We investigated potential correlations between the incidence of each neck use behavior and cervical vertebral morphology for the pooled dataset as well as for each individual cervical region. Consequently, we found a significant, weak correlation between the incidence of total neck use and the morphology of C2 vertebrae (p = 0.034, R^2^ = 0.042), as well as similarly weak correlation between incidence of beak assisted climbing and C2 vertebral morphology (p = 0.043, R^2^ = 0.039) (Supplementary Table 7). Species that have a high incidence of beak assisted climbing rarely occupy distinct areas of morphospace (except for C2), and often coincide with species with minimal or no occurrences of beak assisted climbing (Figure 2, Supplementary Figure 4). These high incidence climbers also do not occupy similar areas of morphospace, often appearing far apart from each other in all studied morphospaces (Figure 2, Supplementary Figure 4). Convergence tests revealed that species with a high incidence of beak assisted climbing and total neck use were not convergent (P >> 0.05). We then visually identified smaller subsets (or pairs) of species with high incidence of neck use behaviors or the same dietary guild that occupied similar areas of phylomorphospace and repeated the convergence tests. None of these further convergence tests were significant (P >> 0.05).

**Figure 2.**
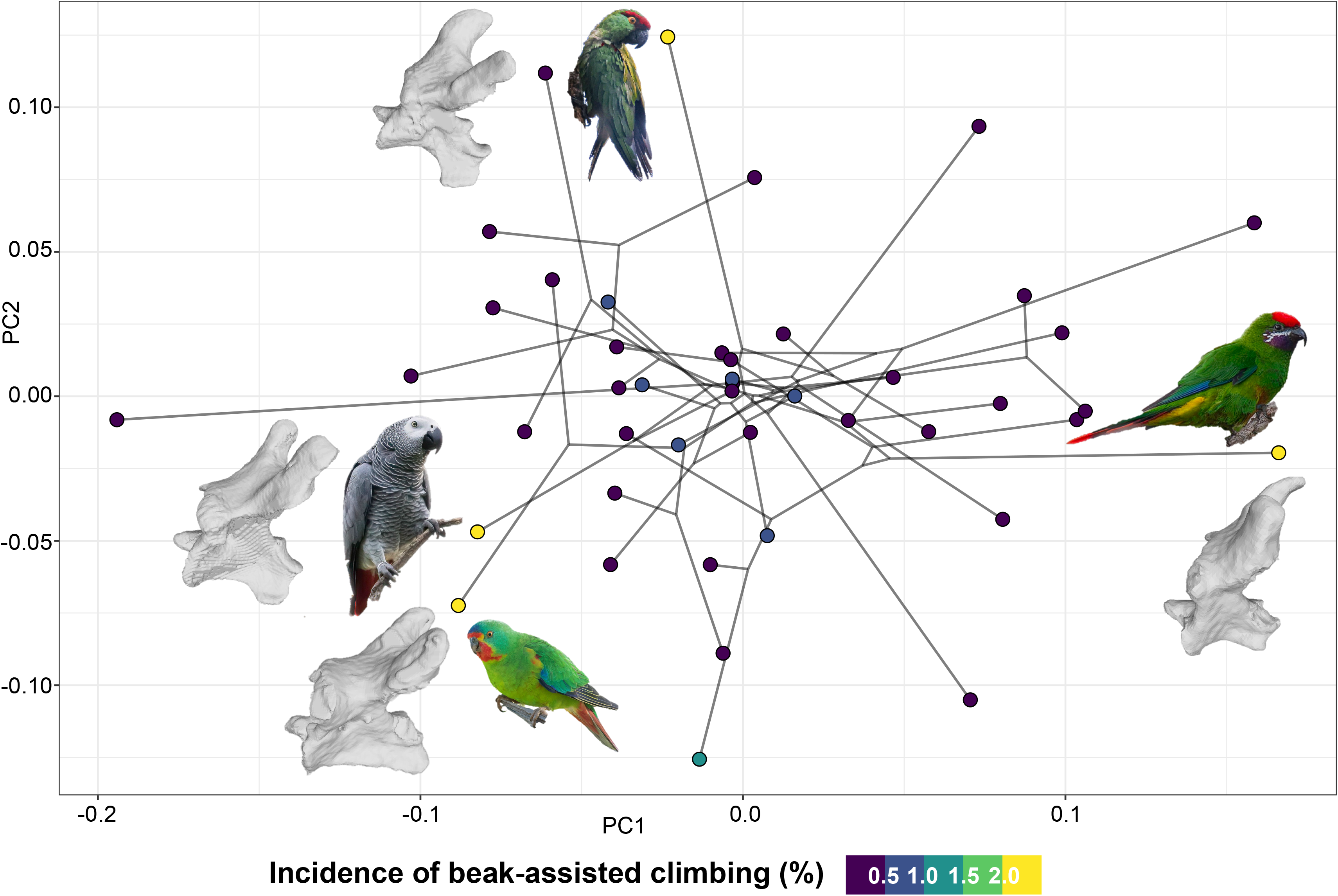
Phylomorphospace of PC1 and PC2 for the C2 vertebrae of parrots. Points are coloured by incidence of beak-assisted climbing. Next to the 4 species of parrot that experience the highest incidence of BAC are photographs of that species as well as the mesh of that species’ C2 vertebrae. Bottom photo: Lathamus discolor (photo credit David Irving, Macaulay Library ID ML613044549). Bottom-left photo: Psittacus erithacus (photo credit Manuel-Fernandez-Bermejo, Macaulay Library ID ML613685886). Top photo: Rhychopsitta pachyrhychna (photo credit Ken Chamberlain, Macaulay Library ID ML608581146). Right photo: Oreopsittacus arfaki (photo credit Robert Tizard, Macaulay Library ID ML613799167).

### The effects of integration on parrot cervical morphology

Parrots display significant integration across all of three of the studied regimes (P< 0.05 for head-neck, neck-forelimb and neck-hindlimb integration, Supplementary Table 8). R PLS values were generally high and ranged from 0.847 for head-neck integration across all vertebrae to 0.522 for neck-forelimb integration in the last cervical vertebrae (Supplementary Table 8). Whereas R PLS values and Z scores seem to decrease towards the proximal end of the cervical column, there are no significant differences between Z scores for integration tests for different vertebrae. Many integrative relationships are not significant when head, forelimb or hindlimb measures are adjusted for body mass (Supplementary Table 8). Only the last cervical vertebrae retains a significant pattern of neck-forelimb integration when adjusted for body mass (Supplementary Table 8). Significance is retained in the pooled dataset, C50% and the last cervical vertebrae for head-neck integration, and no vertebral region retains significance after adjusting hindlimb measurements for body mass (Supplementary Table 8). Across all integration regimes studied, a gradient of body mass can be observed whereby parrots with high body masses displayed extreme values of PLS1 and PLS2 and parrots with low body mass clustered together at the other extreme of PLS1 and PLS2 (Figure 3).

**Figure 3.**
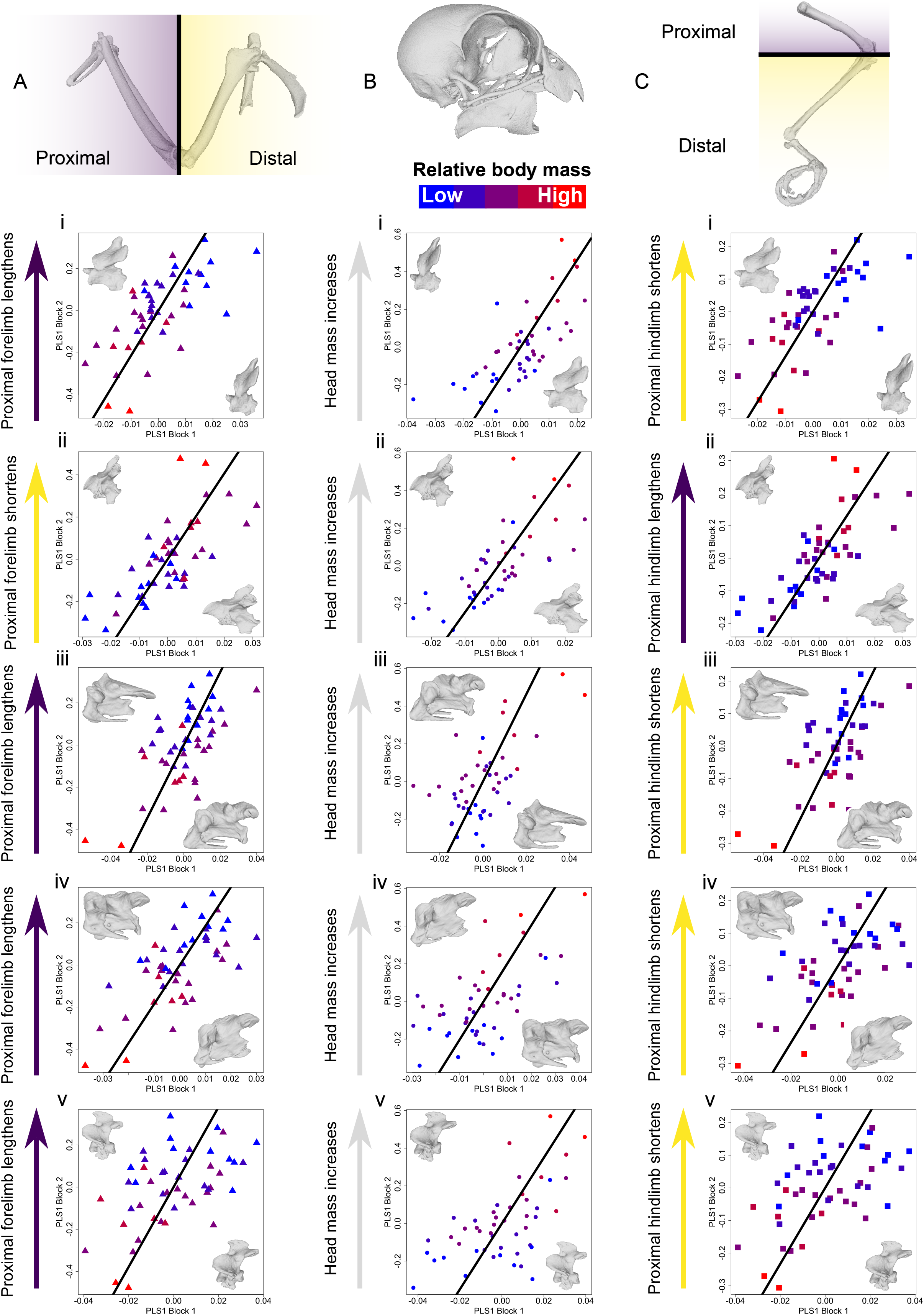
2BPLS plots of vertebral shape versus forelimb proportion (Ai-Av), vertebral shape versus head mass (Bi-Bv) and vertebral shape versus hindlimb proportion (Ci-Cv). Arrows beside the Y-axes indicate what portion of the limb is increasing as PLS2 scores increase: purple denote the proximal portion is lengthening and yellow indicates the distal portion is lengthening. Points within each 2BPLS plot are coloured by body mass. Warped meshes of cervical vertebrae for each plot display shape changes across PLS1. The forelimb is represented by Ara ararauna in A, the skull morphology displayed in B is that of Micropsitta finschii, the hindlimb is represented by Chalcopsitta atra in C.

## Discussion

Here we present the results of the first study to investigate factors that have influenced the morphological variation of psittaciform cervical vertebrae, as well as results from the first study to quantify neck use across Psittaciformes. We find that variation in the morphology of parrot neck vertebrae is governed by a variety of factors including phylogeny, allometry, ecology and integration with both the head and forelimb. Contrary to prior anecdotal evidence, we find that beak-assisted climbing is not a ubiquitous behavior across Psittaciformes and was not observed in 14 of the 48 species studied. Furthermore, we find that a weak but significant relationship exists between the incidence of beak-assisted climbing and the morphological variability of psittaciform C2 vertebrae (Figure 2, Supplementary Table 7). We were also able to qualitatively and quantitatively demonstrate that beak-assisted climbing is not responsible for any convergence in the morphology of cervical vertebrae amongst parrots with similar incidences of this behavior (Figure 2, Supplementary Figure 4). The neck of parrots also displays significant integration with the head, forelimb and hindlimb however these integrative relationships often disappear when body mass is accounted for (Supplementary Table 8).

Evolution of morphological form is often multifactorial in nature due to the interplay of phylogenetic constraints, allometry, ecology and integration with other anatomical components (26–28,51). Indeed, both evolutionary allometry (phylogeny and body size) and phenotypic integration have previously been shown to be some of the factors that influence skull shape variation in parrots (26). The factors that influence morphological variation are therefore broadly similar across the head and neck of Psittaciformes as phylogeny, body mass, and integration all significantly influence the shape variation of the psittaciform neck. A key difference between the skull and neck of parrots is that dietary preference accounts for more morphological variation in the neck of parrots (up to 15% vs 2.4%, Supplementary Table 4). We had expected to observe the opposite pattern as the skull directly manipulates and processes food, not the neck (7,26,52,53). Diet is still a minor (∼15%) component of neck morphological variation however, and the differences observed here may be due to discrepancies in dietary classification schemes between the two studies (26). Allometry accounted for a smaller proportion of cervical shape variation than diet (between 4.4% and 7.5%), however it evidently plays an important role in the integrative relationships between the neck, head and appendicular skeleton as the significance of head-neck, neck-forelimb and neck-hindlimb integration often disappears when body mass is taken into account (Supplementary Table 8). Across many of these integrative relationships we observed that larger parrots clustered together with extreme PLS1 and PLS2 scores (Figure 3) and this clustering may indicate that body mass may be a controlling factor in psittaciform neck integration. Since beak-assisted climbing requires the cooperation of the psittaciform craniocervical and hindlimb skeleton, a coordinated morphological response may be required in order for this multi-body system behavior to occur in larger parrots (16,51,54–56).

Across multiple morphological scales we observe that parrots with different diets and neck use behaviors often have similar neck morphologies (Figures 1 & 2, Supplementary Tables 4-7) and we infer this is a feature of many-to-one mapping (18,19). Although we do observe a significant relationship between diet, neck use and vertebral morphology, we find that these relationships are weak (Supplementary Table 4) and that dietary and neck use groups often heavily overlap in morphospace (Figures 1 & 2). We also find very few significant differences between the pattern of morphological variation across the entire neck among dietary and behavioral groups (Supplementary Table 6). The neck of parrots provides much of the propulsion during beak-assisted climbing. Indeed, kinematic data suggests it is capable of producing relatively greater contractile forces than the human neck (6,57). It has also been anecdotally reported that all parrots engage in this behavior (21). In response to this we hypothesized that beak-assisted climbing would be a highly influential factor in the morphological variance of the cervical column across Psittaciformes, and that this behavior was leading to morphological convergence of cervical vertebrae. In light of the present results, we reject this hypothesis and suggest that form and function are somewhat decoupled in the psittaciform cervical column. Instead of a tight relationship between form and function, we suggest that many-to-one mapping of form to function is allowing parrots to occupy a diverse range of dietary niches and to utilize a wide array of neck use behaviors (18,19), including tripedal locomotion (6,16) and ‘beakiation’ (9). By displaying neither significant morphological convergence or clear morphological adaptations to dietary ecology or neck use we hypothesize that changes to muscle activation patterns and neuromuscular innovation may be responsible for tripedal locomotion and array of neck use behaviors in Psittaciformes, as has previously been suggested (6,16). The neuromuscular pathways associated with neck use behaviors such as beak-assisted climbing must allow for movements of the craniocervical system to be incorporated into the locomotor cycle (6). This may not require a radical neuromuscular innovation as this pathway already exists to accommodate avian head-bobbing (58), and may have been modified by parrots to allow for tripedal locomotion (6,16). Since the neuromuscular pathways associated with the parrot neck are already optimizes for a wide variety of behaviors, these pathways may display plasticity in their ability to adapt to novel functions such as beak-assisted climbing (6,16) and ‘beakiation’ (9).

The findings presented here for parrots may have implications for the mechanisms behind broader avian neck evolution, as the ecological signal in cervical morphological variation is similarly low across Aves (3,20,30). The similarly weak influence of ecology on neck morphology across all birds suggests that many-to-one mapping may be present across Aves. This extrapolation could explain why there is an apparent disconnect between avian cervical form and function: the overall morphological construction of the avian neck is highly conserved and only adapts to behaviors that require specialized kinematic forces, yet the neck still participates in a disparate array of behaviors (3,4). Although further work is required to formally test the presence of many-to-one mapping in the parrot and avian cervical column, it has been shown to be a common feature of organismal design (18,19) that weakens the effect of convergent evolution (59).

## Conclusions

This work represents the first quantification of the presence of tripedal locomotion across parrots and finds that beak-assisted climbing is not a ubiquitous feature of Psittaciformes. We also find that tripedal locomotion, alongside a multitude of other factors, governs a small portion of morphological variability of the parrot neck. Parrots with similar cervical morphologies appear to be able to use their necks to access a wide variety of food types and to both participate and not participate in beak-assisted climbing. This suggests that many-to-one mapping of cervical form to function is a feature of the neck of Psittaciformes, and potentially a feature of neck construction across extant Aves. Without the presence of clear vertebral adaptations to beak-assisted climbing we suggest changes to the neuromuscular control of the cervical column have underpinned the evolution of tripedal locomotion in parrots and this may be a modification of existing neural pathways associated with avian head-bobbing.

## Acknowledgements

This work was funded by a Leverhulme Trust Research Project Grant (award number RPG-2021-088) to RNF. We would like to thank Judith White at the Natural History Museum, Tring for access to specimens.

## Figures & tables

Supplementary Figure 1: Visual representation of the landmark scheme used throughout this study. Mesh is the C25% vertebrae of Pionites melanocephalus.

Supplementary Figure 2: Morphospace of vertebral shape across the neck of all parrots studied. Colours denote vertebral region. Warped meshes of cervical vertebrae display shape change across PC1 and PC2 in anterior (left along PC1, bottom along PC2) view and left lateral (right along PC1, top along PC2).

Supplementary Figure 3: Phenotypic trajectory plot depicting patterns of shape change across the neck of parrots with different dietary niches. Point and line colours denote dietary preference and point shape denotes vertebral region.

Supplementary Figure 4: Phylomorphospace plots for the C25% (A), C50% (B), C75% (C) and last cervical vertebrae (D) of 44 species of parrots. Points are coloured by incidence of beak-assisted climbing.

Supplementary Table 1: Specimen information and metadata for all studied species. Asterisks indicate outlier taxa that were removed from the neck use incidence analysis.

Supplementary Table 2: Landmark scheme used as part of the geometric morphometric component of this analysis.

Supplementary Table 3: Neck-use behaviour data for all species. Asterisks indicate outlier taxa that were removed from the neck use incidence analysis.

Supplementary Table 4: Results from the pMANOVA (phylogenetic multivariate ANOVA) analyses.

Supplementary Table 5: Results summary of post-hoc tests performed after pMANOVA.

Supplementary Table 6: Results summary of the phenotypic trajectory analysis (PTA). MD = magnitude of differences between trajectories, TC = trajectory correlations, SD = trajectory shape differences.

Supplementary Table 7: Behavioural MANOVA results table

Supplementary Table 8: 2BPLS results table

