## Supplementary Information for "The role of many-to-one mapping of vertebral form to function in Psittaciform tripedal locomotion"

**Supplementary Information for Stuart et al. 2024.**

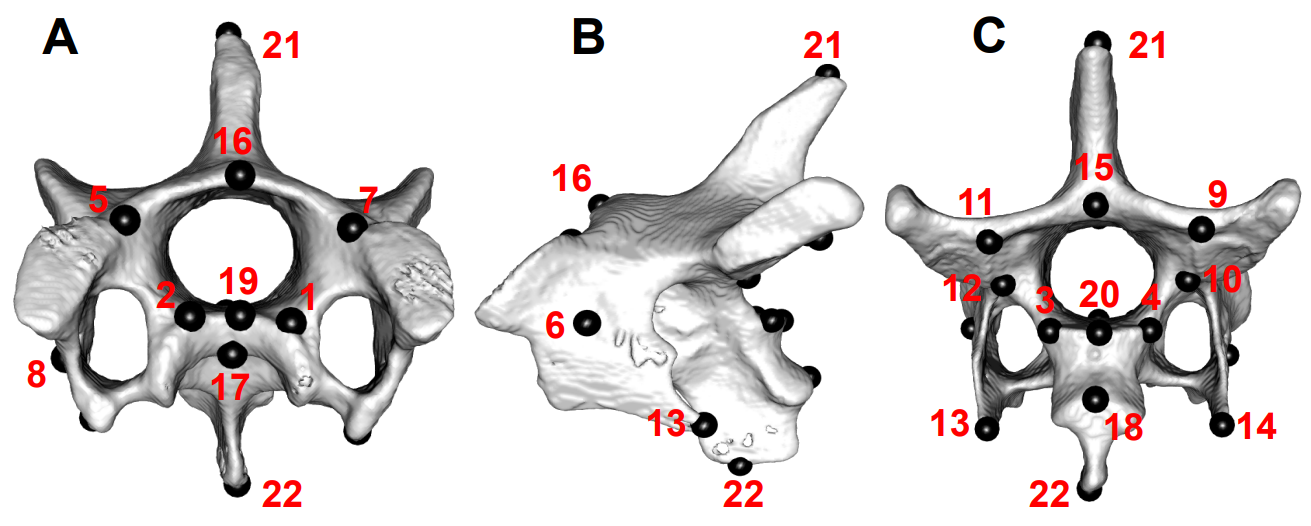
**Supplementary Figure 1:** Visual representation of the landmark scheme used throughout this study. Mesh is the C25% vertebrae of *Pionites melanocephalus*.

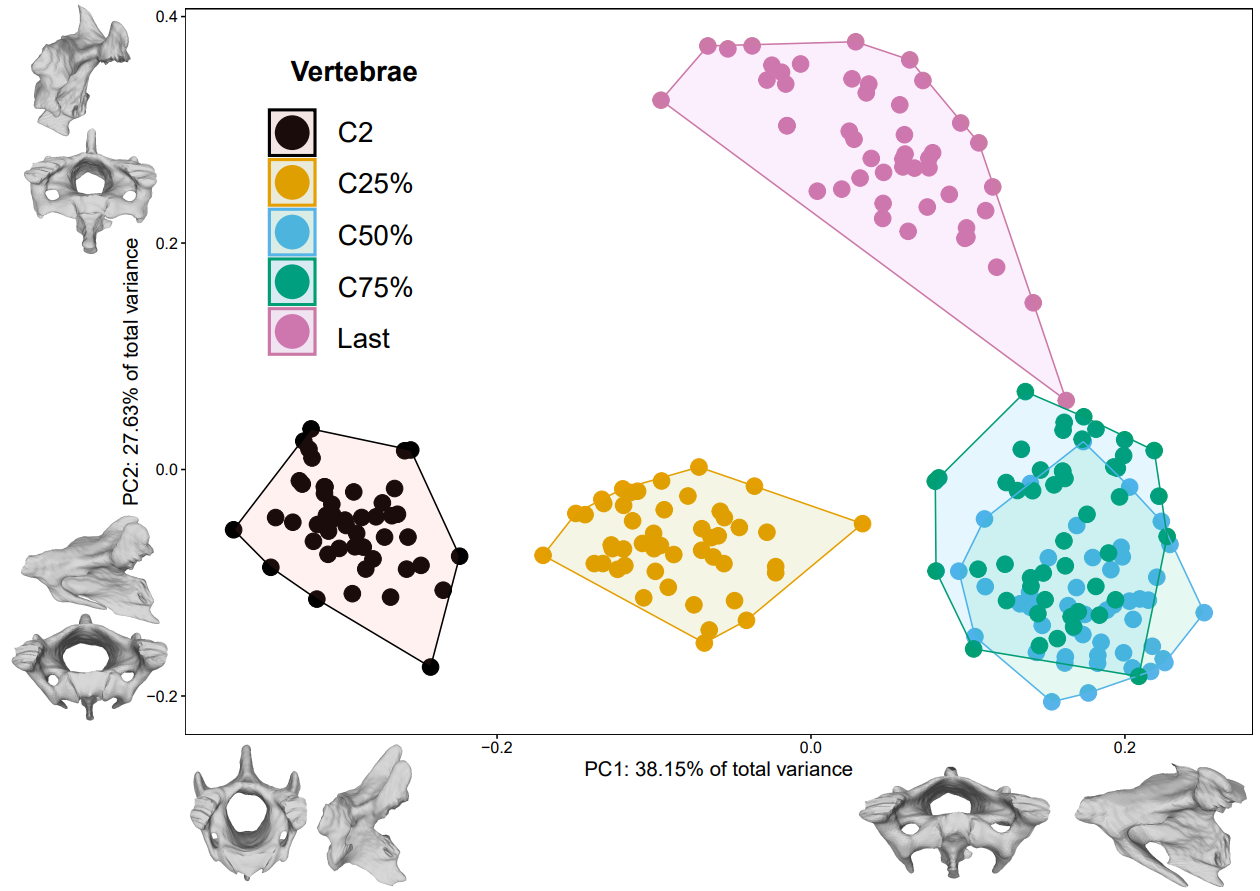
**Supplementary Figure 2:** Morphospace of vertebral shape across the neck of all parrots studied. Colours denote vertebral region. Warped meshes of cervical vertebrae display shape change across PC1 and PC2 in anterior (left along PC1, bottom along PC2) view and left lateral (right along PC1, top along PC2).

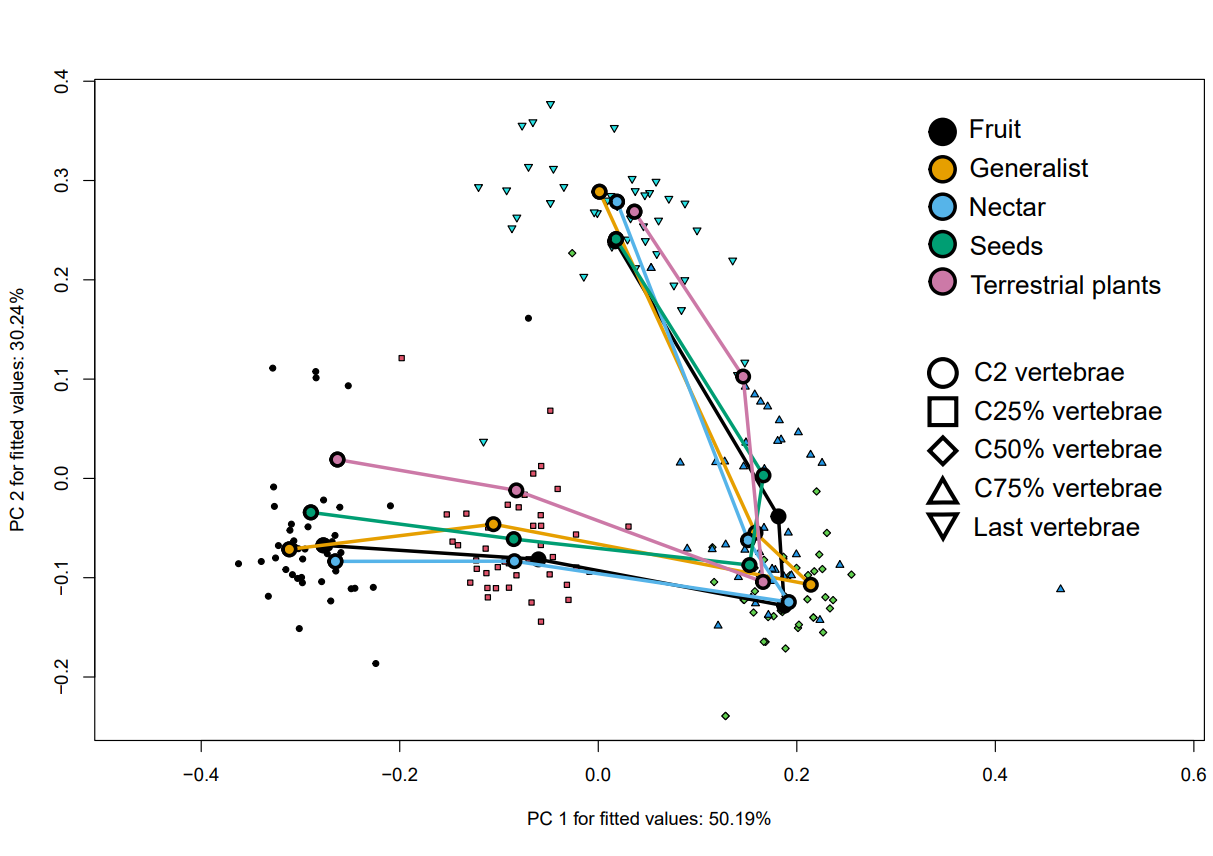
**Supplementary Figure 3:** Phenotypic trajectory plot depicting patterns of shape change across the neck of parrots with different dietary niches. Point and line colours denote dietary preference, whilst point shape denotes vertebral region.

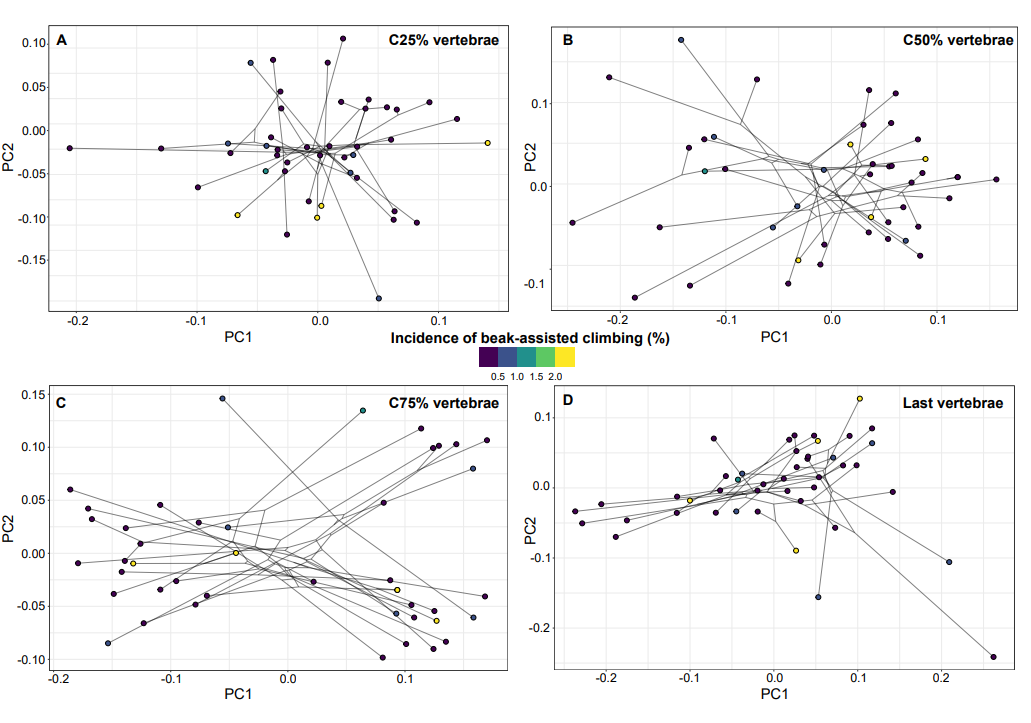
**Supplementary Figure 4:** Phylomorphospace plots for the C25% (A), C50% (B), C75% (C) and last cervical vertebrae (D) of 44 species of parrots. Points are coloured by incidence of beak-assisted climbing.

**Supplementary Table 1:** Specimen information and metadata for all studied species. Asterisks indicate outlier taxa that were removed from the neck use incidence analysis.

| Species | Source | Family | Diet | Foraging | Head mass (kg) | Coracoid (mm) | Scapula (mm) | Humerus (mm) | Radius (mm) | Ulna (mm) | CMC (mm) | Femur (mm) | Tibiotarsus (mm) | Tarsometatarsus (mm) | Body mass (kg) | Head mass (% of body mass) |
| --- | --- | --- | --- | --- | --- | --- | --- | --- | --- | --- | --- | --- | --- | --- | --- | --- |
| Agapornis fischeri | MorphoSource (ID: 000078742) | Psittaculidae | Seeds | ArborealGleaning | 0.006549941 | 19.18 | 20.43 | 20.12 | 21.84 | 24.39 | 16.35 | 21.72 | 28.3 | 12.14 | 0.062561102 | 10.46967012 |
| Agapornis taranta | MorphoSource (ID: 000126120) | Psittaculidae | Seeds | GroundForaging | 0.007671377 | 20.14 | 23.26 | 21.7 | 25.67 | 26.91 | 17.15 | 21.55 | 30.09 | 12.88 | 0.067107264 | 11.43151507 |
| Anodorhynchus hyacinthinus | Scanned at UCL | Psittacidae | Fruit | ArborealGleaning | 0.256912942 | 59.47 | 60.21 | 81.51 | 100.6 | 110.3 | 65.03 | 62.84 | 89.39 | 36.45 | 2.362273234 | 10.87566579 |
| Ara ararauna | Scanned at UCL | Psittacidae | Generalist | ArborealGleaning | 0.170800058 | 56.17 | 57 | 84.65 | 97.69 | 108 | 64.64 | 62.14 | 83.49 | 33.64 | 1.839834307 | 9.283447828 |
| Aratinga holochlora | Scanned at UCL | Psittacidae | Fruit | ArborealGleaning | 0.022613655 | 28.63 | 30.15 | 35.87 | 39.8 | 43.92 | 28.18 | 31.07 | 43.12 | 15.59 | 0.2218498 | 10.19322755 |
| Aratinga leucophthalma | MorphoSource (ID: 000064940) | Psittacidae | Generalist | ArborealGleaning | 0.032620943 | 31.42 | 32.13 | 40.67 | 44.39 | 47.44 | 31.25 | 35.78 | 47.44 | 15.47 | 0.160383337 | 20.33935919 |
| Aratinga wagleri | Scanned at UCL | Psittacidae | Fruit | ArborealGleaning | 0.033329012 | 32.74 | 34.66 | 42.55 | 45.55 | 49.96 | 32.29 | 38.05 | 51.05 | 18.99 | 0.399088074 | 8.351292434 |
| Barnardius zonarius | MorphoSource (ID: 000108655) | Psittaculidae | Fruit | ArborealGleaning | 0.013557705 | 26.94 | 30.88 | 34.54 | 37.18 | 40.71 | 26.77 | 29.58 | 19.61 | 42.64 | 0.118665964 | 11.42509996 |
| Bolbopsittacus lunulatus | MorphoSource (ID: 000078255) | Psittaculidae | Fruit | ArborealGleaning | 0.012838668 | 22.49 | 25.15 | 25.2 | 28.99 | 31.82 | 17.95 | 26.16 | 36.71 | 12.67 | 0.084146108 | 15.25758981 |
| Brotogeris chrysoptera | MorphoSource (ID: 000371680) | Psittacidae | Fruit | ArborealGleaning | 0.008255579 | 20.92 | 24.23 | 28.49 | 32.33 | 34.74 | 20.77 | 24.57 | 35.36 | 12.18 | 0.085669419 | 9.636553039 |
| Chalcopsitta atra | MorphoSource (ID: 000097870) | Psittaculidae | Nectar | ArborealGleaning | 0.023369281 | 31.35 | 34.25 | 37.63 | 37.58 | 42.61 | 28.11 | 36.82 | 51.09 | 18.97 | 0.274544972 | 8.512004729 |
| Charmosyna placentis | MorphoSource (ID: 000077491) | Psittaculidae | TerrestrialPlants | ArborealGleaning | 0.004873685 | 16.3 | 20.21 | 17.31 | 18.71 | 19.83 | 14.74 | 19.4 | 26.32 | 10.69 | 0.047834232 | 10.18869708 |
| Coracopsis nigra | MorphoSource (ID: 000064739) | Psittaculidae | Fruit | ArborealGleaning | 0.021587243 | 32.11 | 31.22 | 57.9 | 49.25 | 53.71 | 36.85 | 38.92 | 56.71 | 20.05 | 0.279134846 | 7.733625274 |
| Diopsittaca nobilis | MorphoSource (ID: 000086019) | Psittacidae | Fruit | ArborealGleaning | 0.030449975 | 29.61 | 30.57 | 36.24 | 38.33 | 42.36 | 26.64 | 24.69 | 40.28 | 14.3 | 0.157077866 | 19.38527418 |
| Enicognathus ferrugineus | MorphoSource (ID: 000113139) | Psittacodae | Generalist | ArborealGleaning | 0.019561738 | 31.22 | 34.67 | 39.13 | 40.8 | 43.91 | 26.27 | 33.91 | 50.79 | 17.67 | 0.205826008 | 9.504016616 |
| Eos squamata | Scanned at UCL | Psittacidae | Generalist | ArborealGleaning | 0.011679357 | 24.43 | 25.45 | 27.17 | 27.48 | 30.86 | 21.54 | 28.19 | 41.38 | 16.72 | 0.159497905 | 7.322576765 |
| Forpus passerinus | MorphoSource (ID: 000064737) | Psittacidae | Seeds | GroundForaging | 0.004219164 | 16.04 | 17.03 | 18.67 | 20.06 | 21.66 | 14.08 | 17.79 | 23.88 | 10.33 | 0.030172672 | 13.98339531 |
| Lathamus discolor | MorphoSource (ID: 000086192) | Psittaculidae | Nectar | ArborealGleaning | 0.00647374 | 20.59 | 23.21 | 22.77 | 22.97 | 25.9 | 19.2 | 22.55 | 33.65 | 12.35 | 0.074634744 | 8.6738959 |
| Loriculus philippensis | MorphoSource (ID: 000064743) | Psittaculidae | Generalist | ArborealGleaning | 0.014509939 | 18.01 | 20.59 | 21.18 | 23.2 | 25.56 | 16.49 | 20.97 | 28.86 | 10.5 | 0.04892724 | 29.65615677 |
| Loriculus vernalis | MorphoSource (ID: 000083832) | Psittaculidae | Fruit | ArborealGleaning | 0.003796745 | 15.66 | 14.81 | 18.16 | 19.76 | 21.42 | 14.07 | 19.63 | 25.84 | 9.28 | 0.040587037 | 9.354575452 |
| Lorius garrulus*** | MorphoSource (ID: 000086194 | Psittaculidae | Nectar | ArborealGleaning | 0.024276751 | 32.38 | 36.05 | 36.51 | 36.78 | 41.53 | 26.66 | 38.13 | 55.3 | 21.13 | 0.201993956 | 12.01855317 |
| Melopsittacus undulatus | MorphoSource (ID: 000098664) | Psittaculidae | Seeds | GroundForaging | 0.003783346 | 18.45 | 20.28 | 18.92 | 19.95 | 21.68 | 15.59 | 19.83 | 25.32 | 11.93 | 0.051157123 | 7.395540988 |
| Micropsitta finschii*** | MorphoSource (ID: 000109367) | Psittaculidae | TerrestrialPlants | ArborealGleaning | 0.002409909 | 13.63 | 14.92 | 14.07 | 14.69 | 16.26 | 10.89 | 14.18 | 20.27 | 7.64 | 0.021527733 | 11.19443928 |
| Myiopsitta monachus | MorphoSource (ID: 000086190) | Psittacidae | Generalist | ArborealGleaning | 0.011484942 | 25.04 | 26.55 | 31.23 | 32.29 | 35.47 | 23.23 | 28.31 | 41.61 | 14.18 | 0.136070186 | 8.440454399 |
| Nannopsittaca panychlora | MorphoSource (ID: 000077517) | Psittacidae | Fruit | ArborealGleaning | 0.005359493 | 17.81 | 22.22 | 19.56 | 20.24 | 21.72 | 16.52 | 21.63 | 28.81 | 10.78 | 0.037224184 | 14.39787908 |
| Neopsephotus bourkii | MorphoSource (ID: 000064729) | Psittaculidae | Seeds | GroundForaging | 0.004609729 | 18.54 | 21.07 | 20.87 | 22.41 | 24.77 | 17.59 | 18.04 | 25.27 | 11.69 | 0.046755768 | 9.859166467 |
| Neopsittacus musschenbroekii | MorphoSource (ID: 000077494) | Psittaculidae | Fruit | ArborealGleaning | 0.007233604 | 18.56 | 20.41 | 21.3 | 22.35 | 24.59 | 16.85 | 21.08 | 34.71 | 13.82 | 0.066446273 | 10.88639539 |
| Nymphicus hollandicus | MorphoSource (ID: 000482459) | Cacatuidae | Seeds | GroundForaging | 0.00809562 | 24.08 | 24.29 | 33.96 | 37.28 | 40.21 | 28.43 | 24.77 | 37.23 | 13.69 | 0.090336492 | 8.9616276 |
| Oreopsittacus arfaki | MorphoSource (ID: 000098663) | Psittaculidae | Generalist | ArborealGleaning | 0.002772029 | 13.64 | 15.71 | 14.7 | 14.96 | 16.73 | 12.08 | 15.06 | 25 | 9.67 | 0.03315034 | 8.361992667 |
| Pionites melanocephalus | MorphoSource (ID: 000126028) | Psittacidae | Seeds | ArborealGleaning | 0.020379079 | 29.52 | 30.36 | 34.01 | 38.06 | 42.52 | 24.99 | 33.62 | 47.04 | 15.97 | 0.191982189 | 10.61508836 |
| Poicephalus meyeri | MorphoSource (ID: 000064733) | Psittacidae | Generalist | ArborealGleaning | 0.01823832 | 26 | 29.72 | 34.37 | 37.33 | 40.4 | 29.99 | 28.4 | 43.34 | 18.33 | 0.15273362 | 11.94126087 |
| Polytelis alexandrae | MorphoSource (ID: 000064730) | Psittaculidae | Seeds | ArborealGleaning | 0.010292031 | 30.7 | 33.07 | 39.6 | 42.76 | 46.52 | 28.59 | 36.66 | 49.34 | 16.51 | 0.150588416 | 6.834543634 |
| Prioniturus discurus | MorphoSource (ID: 000064742) | Psittaculidae | Fruit | ArborealGleaning | 0.018071722 | 43.86 | 44.11 | 75.45 | 88.34 | 94.05 | 45.36 | 56.95 | 82.26 | 26.72 | 0.179887576 | 10.04612014 |
| Probosciger aterrimus | MorphoSource (ID: 000092665) | Cacatuidae | Seeds | ArborealGleaning | 0.14460488 | 20.1 | 19.39 | 26.25 | 28.22 | 28.14 | 20.68 | 29.88 | 44.19 | 15.97 | 0.70870586 | 20.40407568 |
| Psephotus dissimilis | MorphoSource (ID: 000065357) | Psittacidae | Seeds | GroundForaging | 0.005263183 | 20.62 | 23.98 | 25.53 | 28.04 | 30.06 | 20.1 | 21.32 | 30.71 | 13.88 | 0.059305136 | 8.874750574 |
| Pseudeos fuscata*** | MorphoSource (ID: 000086197) | Psittaculidae | Nectar | ArborealGleaning | 0.01169152 | 17.21 | 20.78 | 19.17 | 20.14 | 22.41 | 15.8 | 19.12 | 30.41 | 11.41 | 0.144259928 | 8.104482071 |
| Psilopsiagon aymara | MorphoSource (ID: 000077527) | Psittacidae | Generalist | ArborealGleaning | 0.004737485 | 26.9 | 26.92 | 32.63 | 36.05 | 38.49 | 20.84 | 31.4 | 42.81 | 18.57 | 0.033589269 | 14.10416225 |
| Psittacella brehmii | MorphoSource (ID: 000064940) | Psittaculidae | Generalist | ArborealGleaning | 0.014509939 | 24.74 | 27.61 | 33.07 | 36.13 | 39.24 | 26.97 | 28.92 | 39.71 | 15.53 | 0.122412409 | 11.85332363 |
| Psittacula krameri | MorphoSource (ID: 000064941) | Psittaculidae | Fruit | ArborealGleaning | 0.015500858 | 43.2 | 43.66 | 57.49 | 64.21 | 70.32 | 40.47 | 46.01 | 62.26 | 22.38 | 0.143220743 | 10.82305375 |
| Psittacus erithacus | MorphoSource (ID: 000092678) | Psittacidae | Fruit | ArborealGleaning | 0.027715204 | 30.31 | 27.42 | 38.88 | 43.72 | 48.48 | 30.16 | 33.01 | 46.36 | 15.14 | 0.301151126 | 9.203088286 |
| Pyrilia barrabandi | MorphoSource (ID: 000064736) | Psittacidae | Fruit | ArborealGleaning | 0.018195948 | 25.79 | 29.8 | 35.16 | 37.35 | 40.21 | 25.72 | 30.47 | 40.39 | 14.98 | 0.18587683 | 9.789250226 |
| Pyrrhura frontalis | MorphoSource (ID: 000082499) | Psittacidae | Generalist | ArborealGleaning | 0.010113668 | 20.99 | 22.19 | 25.11 | 27.99 | 30.86 | 19.07 | 20.68 | 29.94 | 12.03 | 0.080408166 | 12.57791155 |
| Pyrrhura picta | MorphoSource (ID: 000064735) | Psittacidae | Fruit | ArborealGleaning | 0.008380642 | 21.14 | 22.19 | 25.28 | 27.97 | 30.89 | 19.57 | 20.48 | 30.37 | 12.09 | 0.075342563 | 11.12338321 |
| Rhynchopsitta pachyrhyncha | MorphoSource (ID: 000064731) | Psittacidae | Seeds | ArborealGleaning | 0.060116907 | 34.17 | 39.1 | 50.94 | 54.72 | 59.87 | 38.95 | 41.29 | 57.08 | 18.37 | 0.460734018 | 13.04807213 |
| Strigops habroptila | MorphoSource (ID: 000110026) | Strigopidae | TerrestrialPlants | GroundForaging | 0.06838413 | 39.79 | 38.18 | 70.14 | 61.79 | 68.91 | 37.17 | 74.12 | 106 | 44.71 | 0.592329286 | 11.5449517 |
| Tanygnathus megalorynchos | MorphoSource (ID: 000108113) | Psittaculidae | Fruit | ArborealGleaning | 0.087737894 | 39.35 | 44.11 | 56.76 | 64.15 | 69.18 | 41.75 | 47.79 | 61.03 | 20.43 | 0.418536256 | 20.96303313 |
| Trichoglossus ornatus*** | MorphoSource (ID: 000359829) | Psittaculidae | Nectar | ArborealGleaning | 0.012640564 | 23.85 | 27.92 | 27.18 | 26.77 | 29.25 | 20.57 | 29.61 | 40.18 | 13.82 | 0.115901126 | 10.90633408 |
| Vini australis | MorphoSource (ID: 000082478) | Psittaculidae | Nectar | ArborealGleaning | 0.005228717 | 17.21 | 20.71 | 20.78 | 21.17 | 22.81 | 16.44 | 25.15 | 22.49 | 25.2 | 0.040587037 | 12.88272657 |

**Supplementary Table 2:** Landmark scheme used as part of the geometric morphometric component of this analysis.

| Landmark number | Description | Presence |
| --- | --- | --- |
| 1 | Maximum curvature of the dorsal and right part of the cranial articular surface | Invariant |
| 2 | Maximum curvature of the dorsal and left part of the cranial articular surface | Invariant |
| 3 | Maximum curvature of the dorsal and left part of the cranial articular surface | Invariant |
| 4 | Maximum curvature of the dorsal and right part of the cranial articular surface | Invariant |
| 5 | Junction of the vertebral arch and left prezygapophyseal facet | Invariant |
| 6 | Lateral-most extent of the left tranverse process | Variant |
| 7 | Junction of the vertebral arch and right prezygapophyseal facet | Invariant |
| 8 | Lateral-most extent of the right transverse process | Variant |
| 9 | Anterior junction of the vertebral arch and right postzygapophyseal facet | Invariant |
| 10 | Posterior junction of the vertebral arch and right postzygapophyseal facet | Invariant |
| 11 | Anterior junction of the vertebral arch and left postzygapophyseal facet | Invariant |
| 12 | Posterior junction of the vertebral arch and left postzygapophyseal facet | Invariant |
| 13 | Caudal-most extent of the left costal process | Variant |
| 14 | Caudal-most extent of the right costal process | Variant |
| 15 | Caudal maximum curvature of vertebral arch | Invariant |
| 16 | Cranial maximum curvature of vertebral arch | Invariant |
| 17 | Middle ventral curvature of the cranial articular surface | Invariant |
| 18 | Middle ventral curvature of the caudal articular surface | Invariant |
| 19 | Middle dorsal curvature of the cranial articular surface | Invariant |
| 20 | Middle dorsal curvature of the caudal articular surface | Invariant |
| 21 | Dorsal-most tip of the neural spine | Invariant |
| 22 | Ventral-most tip of the ventral spine | Invariant |

**Supplementary Table 3:** Neck-use behaviour data for all species. Asterisks indicate outlier taxa that were removed from the neck use incidence analysis.

| Species | Number of photos | Log(Number of photos) | Neck use index | Total number of media | Absolute neck use | Incidences | Number of photos (%) | Total media (%) | Beak assisted climbing | Forceful flexion | Forceful extension | Object carrying | BAC incidences | FF incidence | FE incidences | OC incidences | BAC incidences (%) | FF incidences (%) | FE incidences (%) | OC incidences2 |
| --- | --- | --- | --- | --- | --- | --- | --- | --- | --- | --- | --- | --- | --- | --- | --- | --- | --- | --- | --- | --- |
| Agapornis fischeri | 871 | 2.940018155 | 3 | 914 | 1 | 5 | 0.574052813 | 0.547045952 | 1 | 1 | 1 | 0 | 3 | 1 | 1 | 0 | 0.344431688 | 0.114810563 | 0.114810563 | 0 |
| Agapornis taranta | 328 | 2.515873844 | 1 | 341 | 1 | 1 | 0.304878049 | 0.293255132 | 1 | 0 | 0 | 0 | 1 | 0 | 0 | 0 | 0.304878049 | 0 | 0 | 0 |
| Anodorhynchus hyacinthinus | 1954 | 3.290924559 | 1 | 1977 | 1 | 18 | 0.921187308 | 0.91047041 | 1 | 0 | 0 | 0 | 18 | 0 | 0 | 0 | 0.921187308 | 0 | 0 | 0 |
| Ara ararauna | 2000 | 3.301029996 | 1 | 5852 | 1 | 15 | 0.75 | 0.256322625 | 1 | 0 | 0 | 0 | 15 | 0 | 0 | 0 | 0.75 | 0 | 0 | 0 |
| Aratinga holochlora | 2000 | 3.301029996 | 1 | 3529 | 1 | 9 | 0.45 | 0.255029753 | 1 | 0 | 0 | 0 | 9 | 0 | 0 | 0 | 0.45 | 0 | 0 | 0 |
| Aratinga leucophthalma | 2000 | 3.301029996 | 4 | 6633 | 1 | 21 | 1.05 | 0.316598824 | 1 | 1 | 1 | 1 | 15 | 2 | 3 | 1 | 0.75 | 0.1 | 0.15 | 0.05 |
| Aratinga wagleri | 1697 | 3.229681842 | 1 | 1818 | 1 | 5 | 0.294637596 | 0.275027503 | 1 | 0 | 0 | 0 | 5 | 0 | 0 | 0 | 0.294637596 | 0 | 0 | 0 |
| Barnardius zonarius | 2000 | 3.301029996 | 1 | 4728 | 1 | 4 | 0.2 | 0.084602369 | 1 | 0 | 0 | 0 | 4 | 0 | 0 | 0 | 0.2 | 0 | 0 | 0 |
| Bolbopsittacus lunulatus | 321 | 2.506505032 | 1 | 348 | 1 | 2 | 0.62305296 | 0.574712644 | 1 | 0 | 0 | 0 | 2 | 0 | 0 | 0 | 0.62305296 | 0 | 0 | 0 |
| Brotogeris chrysoptera | 203 | 2.307496038 | 2 | 321 | 1 | 2 | 0.985221675 | 0.62305296 | 1 | 0 | 1 | 0 | 1 | 0 | 1 | 0 | 0.492610837 | 0 | 0.492610837 | 0 |
| Chalcopsitta atra | 23 | 1.361727836 | 0 | 24 | 0 | 0 | 0 | 0 | 0 | 0 | 0 | 0 | 0 | 0 | 0 | 0 | 0 | 0 | 0 | 0 |
| Charmosyna placentis | 91 | 1.959041392 | 0 | 105 | 0 | 0 | 0 | 0 | 0 | 0 | 0 | 0 | 0 | 0 | 0 | 0 | 0 | 0 | 0 | 0 |
| Coracopsis nigra | 302 | 2.480006943 | 1 | 375 | 1 | 1 | 0.331125828 | 0.266666667 | 1 | 0 | 0 | 0 | 1 | 0 | 0 | 0 | 0.331125828 | 0 | 0 | 0 |
| Diopsittaca nobilis | 1495 | 3.174641193 | 2 | 1575 | 1 | 12 | 0.802675585 | 0.761904762 | 1 | 1 | 0 | 0 | 11 | 1 | 0 | 0 | 0.735785953 | 0.066889632 | 0 | 0 |
| Enicognathus ferrugineus | 1588 | 3.200850498 | 3 | 1671 | 1 | 6 | 0.377833753 | 0.359066427 | 1 | 1 | 0 | 1 | 3 | 2 | 0 | 1 | 0.188916877 | 0.125944584 | 0 | 0.062972292 |
| Eos squamata | 97 | 1.986771734 | 1 | 103 | 1 | 1 | 1.030927835 | 0.970873786 | 1 | 0 | 0 | 0 | 1 | 0 | 0 | 0 | 1.030927835 | 0 | 0 | 0 |
| Forpus passerinus | 1735 | 3.239299479 | 2 | 1925 | 1 | 8 | 0.461095101 | 0.415584416 | 1 | 1 | 0 | 0 | 6 | 2 | 0 | 0 | 0.345821326 | 0.115273775 | 0 | 0 |
| Lathamus discolor | 2000 | 3.301029996 | 2 | 2759 | 1 | 46 | 2.3 | 1.66727075 | 1 | 1 | 0 | 0 | 43 | 3 | 0 | 0 | 2.15 | 0.15 | 0 | 0 |
| Loriculus philippensis | 380 | 2.579783597 | 0 | 429 | 0 | 0 | 0 | 0 | 0 | 0 | 0 | 0 | 0 | 0 | 0 | 0 | 0 | 0 | 0 | 0 |
| Loriculus vernalis | 2000 | 3.301029996 | 1 | 4199 | 1 | 8 | 0.4 | 0.190521553 | 1 | 0 | 0 | 0 | 8 | 0 | 0 | 0 | 0.4 | 0 | 0 | 0 |
| Lorius garrulus*** | 59 | 1.770852012 | 1 | 77 | 1 | 3 | 5.084745763 | 3.896103896 | 1 | 0 | 0 | 0 | 3 | 0 | 0 | 0 | 5.084745763 | 0 | 0 | 0 |
| Melopsittacus undulatus | 2000 | 3.301029996 | 2 | 4855 | 1 | 5 | 0.25 | 0.102986612 | 1 | 1 | 0 | 0 | 3 | 2 | 0 | 0 | 0.15 | 0.1 | 0 | 0 |
| Micropsitta finschii*** | 34 | 1.531478917 | 1 | 67 | 1 | 2 | 5.882352941 | 2.985074627 | 1 | 0 | 0 | 0 | 2 | 0 | 0 | 0 | 5.882352941 | 0 | 0 | 0 |
| Myiopsitta monachus | 2000 | 3.301029996 | 3 | 29357 | 1 | 22 | 1.1 | 0.074939537 | 1 | 1 | 0 | 1 | 7 | 3 | 0 | 12 | 0.35 | 0.15 | 0 | 0.6 |
| Nannopsittaca panychlora | 32 | 1.505149978 | 0 | 59 | 0 | 0 | 0 | 0 | 0 | 0 | 0 | 0 | 0 | 0 | 0 | 0 | 0 | 0 | 0 | 0 |
| Neopsephotus bourkii | 670 | 2.826074803 | 0 | 718 | 0 | 0 | 0 | 0 | 0 | 0 | 0 | 0 | 0 | 0 | 0 | 0 | 0 | 0 | 0 | 0 |
| Neopsittacus musschenbroekii | 81 | 1.908485019 | 0 | 89 | 0 | 0 | 0 | 0 | 0 | 0 | 0 | 0 | 0 | 0 | 0 | 0 | 0 | 0 | 0 | 0 |
| Nymphicus hollandicus | 3048 | 3.484014963 | 2 | 3057 | 1 | 9 | 0.295275591 | 0.294406281 | 1 | 1 | 0 | 0 | 5 | 4 | 0 | 0 | 0.164041995 | 0.131233596 | 0 | 0 |
| Oreopsittacus arfaki | 47 | 1.672097858 | 1 | 63 | 1 | 1 | 2.127659574 | 1.587301587 | 1 | 0 | 0 | 0 | 1 | 0 | 0 | 0 | 2.127659574 | 0 | 0 | 0 |
| Pionites melanocephalus | 637 | 2.804139432 | 2 | 792 | 1 | 4 | 0.627943485 | 0.505050505 | 1 | 0 | 0 | 1 | 3 | 0 | 0 | 1 | 0.470957614 | 0 | 0 | 0.156985871 |
| Poicephalus meyeri | 976 | 2.989449818 | 1 | 1061 | 1 | 2 | 0.204918033 | 0.188501414 | 0 | 0 | 0 | 1 | 0 | 0 | 0 | 2 | 0 | 0 | 0 | 0.204918033 |
| Polytelis alexandrae | 53 | 1.72427587 | 0 | 61 | 0 | 0 | 0 | 0 | 0 | 0 | 0 | 0 | 0 | 0 | 0 | 0 | 0 | 0 | 0 | 0 |
| Prioniturus discurus | 55 | 1.740362689 | 0 | 59 | 0 | 0 | 0 | 0 | 0 | 0 | 0 | 0 | 0 | 0 | 0 | 0 | 0 | 0 | 0 | 0 |
| Probosciger aterrimus | 533 | 2.726727209 | 2 | 605 | 1 | 2 | 0.375234522 | 0.330578512 | 1 | 1 | 0 | 0 | 1 | 1 | 0 | 0 | 0.187617261 | 0.187617261 | 0 | 0 |
| Psephotus dissimilis | 803 | 2.904715545 | 1 | 811 | 1 | 3 | 0.373599004 | 0.369913687 | 1 | 0 | 0 | 0 | 3 | 0 | 0 | 0 | 0.373599004 | 0 | 0 | 0 |
| Pseudeos fuscata*** | 18 | 1.255272505 | 1 | 22 | 1 | 2 | 11.11111111 | 9.090909091 | 1 | 0 | 0 | 0 | 2 | 0 | 0 | 0 | 11.11111111 | 0 | 0 | 0 |
| Psilopsiagon aymara | 878 | 2.943494516 | 1 | 906 | 1 | 1 | 0.113895216 | 0.110375276 | 1 | 0 | 0 | 0 | 1 | 0 | 0 | 0 | 0.113895216 | 0 | 0 | 0 |
| Psittacella brehmii | 272 | 2.434568904 | 0 | 259 | 0 | 0 | 0 | 0 | 0 | 0 | 0 | 0 | 0 | 0 | 0 | 0 | 0 | 0 | 0 | 0 |
| Psittacula krameri | 2000 | 3.301029996 | 3 | 32567 | 1 | 11 | 0.55 | 0.033776522 | 1 | 1 | 0 | 1 | 8 | 2 | 0 | 1 | 0.4 | 0.1 | 0 | 0.05 |
| Psittacus erithacus | 434 | 2.63748973 | 2 | 483 | 1 | 13 | 2.995391705 | 2.691511387 | 1 | 1 | 0 | 0 | 9 | 4 | 0 | 0 | 2.073732719 | 0.921658986 | 0 | 0 |
| Pyrilia barrabandi | 568 | 2.754348336 | 3 | 644 | 1 | 15 | 2.64084507 | 2.329192547 | 1 | 1 | 0 | 1 | 4 | 10 | 0 | 1 | 0.704225352 | 1.76056338 | 0 | 0.176056338 |
| Pyrrhura frontalis | 2000 | 3.301029996 | 2 | 3255 | 1 | 5 | 0.25 | 0.153609831 | 1 | 0 | 0 | 1 | 3 | 0 | 0 | 2 | 0.15 | 0 | 0 | 0.1 |
| Pyrrhura picta | 500 | 2.698970004 | 0 | 617 | 0 | 0 | 0 | 0 | 0 | 0 | 0 | 0 | 0 | 0 | 0 | 0 | 0 | 0 | 0 | 0 |
| Rhynchopsitta pachyrhyncha | 135 | 2.130333768 | 2 | 158 | 1 | 4 | 2.962962963 | 2.53164557 | 1 | 0 | 1 | 0 | 3 | 0 | 1 | 0 | 2.222222222 | 0 | 0.740740741 | 0 |
| Strigops habroptila | 49 | 1.69019608 | 0 | 66 | 0 | 0 | 0 | 0 | 0 | 0 | 0 | 0 | 0 | 0 | 0 | 0 | 0 | 0 | 0 | 0 |
| Tanygnathus megalorynchos | 94 | 1.973127854 | 0 | 104 | 0 | 0 | 0 | 0 | 0 | 0 | 0 | 0 | 0 | 0 | 0 | 0 | 0 | 0 | 0 | 0 |
| Trichoglossus ornatus*** | 40 | 1.602059991 | 1 | 51 | 1 | 2 | 5 | 3.921568627 | 1 | 0 | 0 | 0 | 2 | 0 | 0 | 0 | 5 | 0 | 0 | 0 |
| Vini australis | 33 | 1.51851394 | 0 | 42 | 0 | 0 | 0 | 0 | 0 | 0 | 0 | 0 | 0 | 0 | 0 | 0 | 0 | 0 | 0 | 0 |

**Supplementary Table 4:** Results from the pMANOVA (phylogenetic multivariate ANOVA) analyses.

| Vertebrae | Dependent variable | SS | MS | R^2^ | F | Z | P-value |
| --- | --- | --- | --- | --- | --- | --- | --- |
| All | Body mass | 0.014696826 | 0.014696826 | 0.045302134 | 1.898072084 | 2.649880467 | 0.004 |
| All | Diet | 0.049039176 | 0.012259794 | 0.122328982 | 1.498325152 | 2.63787287 | 0.004 |
| All | Foraging | 0.006170808 | 0.006170808 | 0.015393177 | 0.719156247 | -0.703963161 | 0.758 |
| All | Head mass | 0.011350438 | 0.011350438 | 0.034987082 | 1.450222309 | 1.492319113 | 0.074 |
| All | Head mass (%) | 0.012568544 | 0.012568544 | 0.038741825 | 1.612129835 | 1.588185253 | 0.07 |
| C2 | Body mass | 0.004615247 | 0.004615247 | 0.075037293 | 3.24498675 | 4.086707234 | 0.001 |
| C2 | Diet | 0.010464559 | 0.00261614 | 0.139570013 | 1.743753311 | 3.039476357 | 0.004 |
| C2 | Foraging | 0.000782472 | 0.000782472 | 0.01043614 | 0.485125286 | -1.381718049 | 0.917 |
| C2 | Head mass | 0.002163301 | 0.002163301 | 0.035172169 | 1.458173882 | 1.247212491 | 0.11 |
| C2 | Head mass (%) | 0.001678403 | 0.001678403 | 0.027288429 | 1.122159124 | 0.578399726 | 0.276 |
| C25% | Body mass | 0.003415743 | 0.003415743 | 0.059744989 | 2.541650432 | 3.299448931 | 0.002 |
| C25% | Diet | 0.00773899 | 0.001934748 | 0.120919982 | 1.478693379 | 2.238690437 | 0.014 |
| C25% | Foraging | 0.000662174 | 0.000662174 | 0.010346328 | 0.480906695 | -1.58984496 | 0.934 |
| C25% | Head mass | 0.002361857 | 0.002361857 | 0.041311396 | 1.723662739 | 1.837323157 | 0.033 |
| C25% | Head mass (%) | 0.002050096 | 0.002050096 | 0.03585837 | 1.48768062 | 1.25992699 | 0.103 |
| C50% | Body mass | 0.001363533 | 0.001363533 | 0.022882571 | 0.936737801 | 0.214901449 | 0.419 |
| C50% | Diet | 0.009324703 | 0.002331176 | 0.109909829 | 1.327428046 | 1.3105415 | 0.093 |
| C50% | Foraging | 0.001369229 | 0.001369229 | 0.016139032 | 0.754573566 | -0.197734168 | 0.578 |
| C50% | Head mass | 0.001626827 | 0.001626827 | 0.027301118 | 1.122695561 | 0.659540855 | 0.261 |
| C50% | Head mass (%) | 0.000633362 | 0.000633362 | 0.010628973 | 0.429726471 | -1.457274788 | 0.928 |
| C75% | Body mass | 0.002406797 | 0.002406797 | 0.029462514 | 1.21427619 | 0.708421836 | 0.24 |
| C75% | Diet | 0.015304971 | 0.003826243 | 0.151422513 | 1.91825973 | 2.275815241 | 0.014 |
| C75% | Foraging | 0.002429678 | 0.002429678 | 0.024038462 | 1.133004954 | 0.600238458 | 0.28 |
| C75% | Head mass | 0.001654707 | 0.001654707 | 0.020255898 | 0.826987298 | 0.119893481 | 0.449 |
| C75% | Head mass (%) | 0.005125262 | 0.005125262 | 0.062740289 | 2.677605309 | 1.921493373 | 0.026 |
| Last | Body mass | 0.002895506 | 0.002895506 | 0.044918406 | 1.881238494 | 1.780842221 | 0.035 |
| Last | Diet | 0.006205953 | 0.001551488 | 0.081671001 | 0.956044364 | 0.007421236 | 0.5 |
| Last | Foraging | 0.000927255 | 0.000927255 | 0.012202781 | 0.56826229 | -0.839971271 | 0.804 |
| Last | Head mass | 0.003543746 | 0.003543746 | 0.054974645 | 2.326906658 | 2.133041214 | 0.014 |
| Last | Head mass (%) | 0.00308142 | 0.00308142 | 0.047802504 | 2.008091986 | 1.711601041 | 0.04 |
| All | BAC incidence (%) | 0.009940858 | 0.009940858 | 0.02899259 | 1.254046866 | 1.037100032 | 0.152 |
| All | Forceful flexion incidence (%) | 0.007152234 | 0.007152234 | 0.020859547 | 0.894765387 | -0.265613232 | 0.588 |
| All | Forceful extension incidence (%) | 0.00717818 | 0.00717818 | 0.02093522 | 0.898080747 | 0.095586659 | 0.476 |
| All | Object carrying incidence (%) | 0.005366669 | 0.005366669 | 0.015651931 | 0.667834015 | -0.805611547 | 0.789 |
| All | Total neck use incidence (%) | 0.010709176 | 0.010709176 | 0.031233396 | 1.354095631 | 1.098667656 | 0.138 |
| C2 | BAC incidence (%) | 0.002300112 | 0.002300112 | 0.038507923 | 1.6821072 | 1.772081961 | 0.043 |
| C2 | Forceful flexion incidence (%) | 0.001860842 | 0.001860842 | 0.031153765 | 1.350532315 | 0.668598356 | 0.243 |
| C2 | Forceful extension incidence (%) | 0.001640481 | 0.001640481 | 0.027464535 | 1.186085743 | 0.828177279 | 0.205 |
| C2 | Object carrying incidence (%) | 0.001150412 | 0.001150412 | 0.019259925 | 0.824802505 | -0.057571537 | 0.512 |
| C2 | Total neck use incidence (%) | 0.002537638 | 0.002537638 | 0.042484526 | 1.86352089 | 1.876474792 | 0.034 |
| C25% | BAC incidence (%) | 0.001648793 | 0.001648793 | 0.030337302 | 1.314030834 | 1.166253064 | 0.138 |
| C25% | Forceful flexion incidence (%) | 0.001328287 | 0.001328287 | 0.024440085 | 1.052199429 | 0.104243541 | 0.463 |
| C25% | Forceful extension incidence (%) | 0.001467075 | 0.001467075 | 0.026993739 | 1.165189884 | 0.808802068 | 0.233 |
| C25% | Object carrying incidence (%) | 0.000900542 | 0.000900542 | 0.016569707 | 0.70765331 | -0.479485522 | 0.67 |
| C25% | Total neck use incidence (%) | 0.001699124 | 0.001699124 | 0.031263379 | 1.355437459 | 1.021459223 | 0.156 |
| C50% | BAC incidence (%) | 0.001494815 | 0.001494815 | 0.020781728 | 0.891356508 | 0.123142618 | 0.444 |
| C50% | Forceful flexion incidence (%) | 0.001044366 | 0.001044366 | 0.014519339 | 0.618796756 | -0.803729711 | 0.806 |
| C50% | Forceful extension incidence (%) | 0.001540309 | 0.001540309 | 0.021414207 | 0.919078019 | 0.261469532 | 0.386 |
| C50% | Object carrying incidence (%) | 0.000916118 | 0.000916118 | 0.012736366 | 0.541828285 | -0.697288062 | 0.777 |
| C50% | Total neck use incidence (%) | 0.001490165 | 0.001490165 | 0.020717076 | 0.888524819 | -0.018697047 | 0.506 |
| C75% | BAC incidence (%) | 0.003537238 | 0.003537238 | 0.040028277 | 1.751288719 | 1.259349701 | 0.101 |
| C75% | Forceful flexion incidence (%) | 0.001481473 | 0.001481473 | 0.016764715 | 0.71612365 | -0.350028187 | 0.631 |
| C75% | Forceful extension incidence (%) | 0.00175822 | 0.00175822 | 0.019896461 | 0.852615391 | 0.079274353 | 0.488 |
| C75% | Object carrying incidence (%) | 0.001067012 | 0.001067012 | 0.01207458 | 0.513330605 | -0.823796323 | 0.78 |
| C75% | Total neck use incidence (%) | 0.003882376 | 0.003882376 | 0.043933935 | 1.930018577 | 1.359212155 | 0.091 |
| Last | BAC incidence (%) | 0.000959899 | 0.000959899 | 0.014013445 | 0.596929723 | -0.803654198 | 0.783 |
| Last | Forceful flexion incidence (%) | 0.001437267 | 0.001437267 | 0.02098248 | 0.900151556 | -0.11480198 | 0.554 |
| Last | Forceful extension incidence (%) | 0.000772095 | 0.000772095 | 0.011271725 | 0.47880948 | -0.947055738 | 0.818 |
| Last | Object carrying incidence (%) | 0.001332584 | 0.001332584 | 0.019454227 | 0.833288527 | -0.017576083 | 0.489 |
| Last | Total neck use incidence (%) | 0.001099873 | 0.001099873 | 0.016056911 | 0.685395598 | -0.6133316 | 0.728 |

**Supplementary Table 5:** Results summary of post-hoc tests performed after pMANOVA.

| Vertebrae | Comparison | P-value |
| --- | --- | --- |
| C2 | Fruit:TerrestrialPlants | 0.043 |
| C2 | Generalist:TerrestrialPlants | 0.04 |
| C2 | Nectar:Seeds | 0.026 |
| C2 | Nectar:TerrestrialPlants | 0.012 |
| C25% | Fruit:TerrestrialPlants | 0.049 |
| C25% | Generalist:TerrestrialPlants | 0.04 |
| C25% | Nectar:Seeds | 0.044 |
| C25% | Nectar:TerrestrialPlants | 0.015 |
| C75% | Generalist:Seeds | 0.032 |
| C75% | Generalist:TerrestrialPlants | 0.017 |
| C75% | Nectar:TerrestrialPlants | 0.02 |

**Supplementary Table 6:** Results summary of the phenotypic trajectory analysis (PTA). MD = magnitude of differences between trajectories, TC = trajectory correlations, SD = trajectory shape differences.

| Trajectory metric | Dependent variable | Comparison | d | UCL (95%) | Z | P-value | r (TC only) | Angle (TC only) |
| --- | --- | --- | --- | --- | --- | --- | --- | --- |
| MD | Diet | Fruit:Generalist | 0.168698 | 0.13599728 | 1.891248 | 0.035 |  |  |
| MD | Diet | Fruit:Nectar | 0.03768 | 0.21628403 | -0.61002 | 0.685 |  |  |
| MD | Diet | Fruit:Seeds | 0.067878 | 0.14427202 | 0.478354 | 0.3 |  |  |
| MD | Diet | Fruit:TerrestrialPlants | 0.269028 | 0.39938042 | 0.811423 | 0.195 |  |  |
| MD | Diet | Generalist:Nectar | 0.131019 | 0.20192104 | 0.964297 | 0.155 |  |  |
| MD | Diet | Generalist:Seeds | 0.236576 | 0.13830749 | 2.547868 | 0.005 |  |  |
| MD | Diet | Generalist:TerrestrialPlants | 0.10033 | 0.38894405 | -0.27124 | 0.62 |  |  |
| MD | Diet | Nectar:Seeds | 0.105557 | 0.20080649 | 0.665956 | 0.265 |  |  |
| MD | Diet | Nectar:TerrestrialPlants | 0.231348 | 0.37230044 | 0.89983 | 0.22 |  |  |
| MD | Diet | Seeds:TerrestrialPlants | 0.336906 | 0.376187 | 1.308306 | 0.095 |  |  |
| MD | Foraging | ArborealGleaning:GroundForaging | 0.054717 | 0.27403751 | -0.78913 | 0.74 |  |  |
| MD | Family | Cacatuidae:Psittacidae | 0.081051 | 0.61070303 | -1.2213 | 0.9 |  |  |
| MD | Family | Cacatuidae:Psittaculidae | 0.010209 | 0.61085032 | -2.49847 | 0.995 |  |  |
| MD | Family | Psittacidae:Psittaculidae | 0.09126 | 0.11647984 | 1.030646 | 0.16 |  |  |
| MD | Beak-assisted climbing | No:Yes | 0.047763 | 0.07320157 | 0.930977 | 0.145 |  |  |
| MD | Neck Use Index | 0:1 | 0.071216 | 0.08615863 | 1.310802 | 0.105 |  |  |
| MD | Neck Use Index | 0:2 | 0.006205 | 0.08977375 | -1.27728 | 0.875 |  |  |
| MD | Neck Use Index | 0:3 | 0.106746 | 0.1515518 | 0.935258 | 0.185 |  |  |
| MD | Neck Use Index | 1:2 | 0.065011 | 0.09171607 | 0.926707 | 0.19 |  |  |
| MD | Neck Use Index | 1:3 | 0.177962 | 0.16061446 | 1.790643 | 0.03 |  |  |
| MD | Neck Use Index | 2:3 | 0.112951 | 0.16062287 | 1.074163 | 0.16 |  |  |
| MD | Forceful Flexion | No:Yes | 0.052178 | 0.08257947 | 0.726994 | 0.25 |  |  |
| MD | Forceful Extension | No:Yes | 0.119035 | 0.18914724 | 0.547924 | 0.3 |  |  |
| MD | Object Carrying | No:Yes | 0.103034 | 0.11330305 | 1.249967 | 0.085 |  |  |
| TC | Diet | Fruit:Generalist |  | 0.54506823 | -1.61562 | 0.945 | 0.98108573 | 0.194803474 |
| TC | Diet | Fruit:Nectar |  | 0.60961857 | -0.01266 | 0.51 | 0.94515128 | 0.332738881 |
| TC | Diet | Fruit:Seeds |  | 0.46209771 | 0.699278 | 0.23 | 0.93676783 | 0.357519299 |
| TC | Diet | Fruit:TerrestrialPlants |  | 0.97734961 | 1.528632 | 0.065 | 0.6067945 | 0.918774767 |
| TC | Diet | Generalist:Nectar |  | 0.67220318 | -0.15508 | 0.585 | 0.93835524 | 0.352955311 |
| TC | Diet | Generalist:Seeds |  | 0.592066 | 0.077254 | 0.475 | 0.94490939 | 0.333478647 |
| TC | Diet | Generalist:TerrestrialPlants |  | 1.02889937 | 1.341909 | 0.1 | 0.64860658 | 0.865044056 |
| TC | Diet | Nectar:Seeds |  | 0.60724797 | 0.483967 | 0.305 | 0.91907642 | 0.405065919 |
| TC | Diet | Nectar:TerrestrialPlants |  | 1.12950611 | 1.236809 | 0.105 | 0.64064999 | 0.875451837 |
| TC | Diet | Seeds:TerrestrialPlants |  | 1.03175154 | 0.86771 | 0.19 | 0.77796977 | 0.679368302 |
| TC | Foraging | ArborealGleaning:GroundForaging |  | 0.65281544 | 0.735771 | 0.225 | 0.91725681 | 0.40965871 |
| TC | Family | Cacatuidae:Psittacidae |  | 1.26288775 | 0.393943 | 0.35 | 0.79136659 | 0.657755164 |
| TC | Family | Cacatuidae:Psittaculidae |  | 1.33096585 | 0.43432 | 0.355 | 0.78279298 | 0.671654809 |
| TC | Family | Psittacidae:Psittaculidae |  | 0.40716499 | -0.90742 | 0.825 | 0.98038848 | 0.198373175 |
| TC | Beak-assisted climbing | No:Yes |  | 0.29122192 | 0.640875 | 0.275 | 0.97832393 | 0.208589701 |
| TC | Neck Use Index | 0:1 |  | 0.38385352 | 0.286684 | 0.4 | 0.97760597 | 0.212028836 |
| TC | Neck Use Index | 0:2 |  | 0.48519905 | 0.180689 | 0.455 | 0.97417569 | 0.22775525 |
| TC | Neck Use Index | 0:3 |  | 0.56310432 | -1.49131 | 0.945 | 0.9847644 | 0.174782427 |
| TC | Neck Use Index | 1:2 |  | 0.42485468 | -0.51429 | 0.71 | 0.98455316 | 0.175992995 |
| TC | Neck Use Index | 1:3 |  | 0.52172256 | 0.006631 | 0.52 | 0.96933839 | 0.248272417 |
| TC | Neck Use Index | 2:3 |  | 0.56542157 | -0.00764 | 0.485 | 0.96202368 | 0.276474768 |
| TC | Forceful Flexion | No:Yes |  | 0.36901479 | 0.013955 | 0.48 | 0.98592275 | 0.167990493 |
| TC | Forceful Extension | No:Yes |  | 0.65800837 | -0.23263 | 0.585 | 0.96898096 | 0.249722825 |
| TC | Object Carrying | No:Yes |  | 0.41235212 | -1.119 | 0.89 | 0.98944248 | 0.145438316 |
| SD | Diet | Fruit:Generalist | 0.117977 | 0.19683688 | -0.39444 | 0.66 |  |  |
| SD | Diet | Fruit:Nectar | 0.133156 | 0.2207682 | -0.34047 | 0.655 |  |  |
| SD | Diet | Fruit:Seeds | 0.070452 | 0.1750505 | -1.71166 | 0.965 |  |  |
| SD | Diet | Fruit:TerrestrialPlants | 0.257887 | 0.29762322 | 0.925408 | 0.165 |  |  |
| SD | Diet | Generalist:Nectar | 0.115772 | 0.23365787 | -1.07451 | 0.87 |  |  |
| SD | Diet | Generalist:Seeds | 0.16584 | 0.1926383 | 0.931249 | 0.2 |  |  |
| SD | Diet | Generalist:TerrestrialPlants | 0.328428 | 0.30808699 | 1.839497 | 0.02 |  |  |
| SD | Diet | Nectar:Seeds | 0.175064 | 0.22169692 | 0.529716 | 0.29 |  |  |
| SD | Diet | Nectar:TerrestrialPlants | 0.327859 | 0.31473716 | 1.800568 | 0.035 |  |  |
| SD | Diet | Seeds:TerrestrialPlants | 0.21117 | 0.30671097 | 0.18428 | 0.445 |  |  |
| SD | Foraging | ArborealGleaning:GroundForaging | 0.170199 | 0.21237057 | 0.619746 | 0.27 |  |  |
| SD | Family | Cacatuidae:Psittacidae | 0.199084 | 0.38644685 | -0.46739 | 0.645 |  |  |
| SD | Family | Cacatuidae:Psittaculidae | 0.251457 | 0.38559546 | 0.197987 | 0.41 |  |  |
| SD | Family | Psittacidae:Psittaculidae | 0.093424 | 0.15749292 | -0.55258 | 0.69 |  |  |
| SD | Beak-assisted climbing | No:Yes | 0.053611 | 0.10317476 | -0.93741 | 0.84 |  |  |
| SD | Neck Use Index | 0:1 | 0.062727 | 0.11613426 | -0.8651 | 0.825 |  |  |
| SD | Neck Use Index | 0:2 | 0.091752 | 0.12782664 | 0.26241 | 0.39 |  |  |
| SD | Neck Use Index | 0:3 | 0.105314 | 0.1673802 | -0.22997 | 0.595 |  |  |
| SD | Neck Use Index | 1:2 | 0.086362 | 0.12593741 | 0.072924 | 0.47 |  |  |
| SD | Neck Use Index | 1:3 | 0.122724 | 0.15883321 | 0.34834 | 0.365 |  |  |
| SD | Neck Use Index | 2:3 | 0.13937 | 0.16936133 | 0.832677 | 0.21 |  |  |
| SD | Forceful Flexion | No:Yes | 0.049302 | 0.10499797 | -1.44539 | 0.945 |  |  |
| SD | Forceful Extension | No:Yes | 0.080582 | 0.16865148 | -1.18456 | 0.885 |  |  |
| SD | Object Carrying | No:Yes | 0.063768 | 0.12990949 | -1.11643 | 0.895 |  |  |

**Supplementary Table 7:** Behavioural MANOVA results table

| Vertebrae | Dependent variable | SS | MS | R^2^ | F | Z | P-value |
| --- | --- | --- | --- | --- | --- | --- | --- |
| All | BAC incidence (%) | 0.009941 | 0.009941 | 0.028993 | 1.254047 | 1.0371 | 0.152 |
| All | Forceful flexion incidence (%) | 0.007152 | 0.007152 | 0.02086 | 0.894765 | -0.26561 | 0.588 |
| All | Forceful extension incidence (%) | 0.007178 | 0.007178 | 0.020935 | 0.898081 | 0.095587 | 0.476 |
| All | Object carrying incidence (%) | 0.005367 | 0.005367 | 0.015652 | 0.667834 | -0.80561 | 0.789 |
| All | Total neck use incidence (%) | 0.010709 | 0.010709 | 0.031233 | 1.354096 | 1.098668 | 0.138 |
| C2 | BAC incidence (%) | 0.0023 | 0.0023 | 0.038508 | 1.682107 | 1.772082 | 0.043 |
| C2 | Forceful flexion incidence (%) | 0.001861 | 0.001861 | 0.031154 | 1.350532 | 0.668598 | 0.243 |
| C2 | Forceful extension incidence (%) | 0.00164 | 0.00164 | 0.027465 | 1.186086 | 0.828177 | 0.205 |
| C2 | Object carrying incidence (%) | 0.00115 | 0.00115 | 0.01926 | 0.824803 | -0.05757 | 0.512 |
| C2 | Total neck use incidence (%) | 0.002538 | 0.002538 | 0.042485 | 1.863521 | 1.876475 | 0.034 |
| C25% | BAC incidence (%) | 0.001649 | 0.001649 | 0.030337 | 1.314031 | 1.166253 | 0.138 |
| C25% | Forceful flexion incidence (%) | 0.001328 | 0.001328 | 0.02444 | 1.052199 | 0.104244 | 0.463 |
| C25% | Forceful extension incidence (%) | 0.001467 | 0.001467 | 0.026994 | 1.16519 | 0.808802 | 0.233 |
| C25% | Object carrying incidence (%) | 0.000901 | 0.000901 | 0.01657 | 0.707653 | -0.47949 | 0.67 |
| C25% | Total neck use incidence (%) | 0.001699 | 0.001699 | 0.031263 | 1.355437 | 1.021459 | 0.156 |
| C50% | BAC incidence (%) | 0.001495 | 0.001495 | 0.020782 | 0.891357 | 0.123143 | 0.444 |
| C50% | Forceful flexion incidence (%) | 0.001044 | 0.001044 | 0.014519 | 0.618797 | -0.80373 | 0.806 |
| C50% | Forceful extension incidence (%) | 0.00154 | 0.00154 | 0.021414 | 0.919078 | 0.26147 | 0.386 |
| C50% | Object carrying incidence (%) | 0.000916 | 0.000916 | 0.012736 | 0.541828 | -0.69729 | 0.777 |
| C50% | Total neck use incidence (%) | 0.00149 | 0.00149 | 0.020717 | 0.888525 | -0.0187 | 0.506 |
| C75% | BAC incidence (%) | 0.003537 | 0.003537 | 0.040028 | 1.751289 | 1.25935 | 0.101 |
| C75% | Forceful flexion incidence (%) | 0.001481 | 0.001481 | 0.016765 | 0.716124 | -0.35003 | 0.631 |
| C75% | Forceful extension incidence (%) | 0.001758 | 0.001758 | 0.019896 | 0.852615 | 0.079274 | 0.488 |
| C75% | Object carrying incidence (%) | 0.001067 | 0.001067 | 0.012075 | 0.513331 | -0.8238 | 0.78 |
| C75% | Total neck use incidence (%) | 0.003882 | 0.003882 | 0.043934 | 1.930019 | 1.359212 | 0.091 |
| Last | BAC incidence (%) | 0.00096 | 0.00096 | 0.014013 | 0.59693 | -0.80365 | 0.783 |
| Last | Forceful flexion incidence (%) | 0.001437 | 0.001437 | 0.020982 | 0.900152 | -0.1148 | 0.554 |
| Last | Forceful extension incidence (%) | 0.000772 | 0.000772 | 0.011272 | 0.478809 | -0.94706 | 0.818 |
| Last | Object carrying incidence (%) | 0.001333 | 0.001333 | 0.019454 | 0.833289 | -0.01758 | 0.489 |
| Last | Total neck use incidence (%) | 0.0011 | 0.0011 | 0.016057 | 0.685396 | -0.61333 | 0.728 |

**Supplementary Table 8:** 2BPLS results table

| Vertebrae | Dependent variable | Z | p | R PLS |
| --- | --- | --- | --- | --- |
| All | Forelimb | 3.631169 | 0.001 | 0.839626 |
| C2 | Forelimb | 2.655752 | 0.004 | 0.671065 |
| C25% | Forelimb | 2.714471 | 0.002 | 0.70859 |
| C50% | Forelimb | 3.25768 | 0.001 | 0.64909 |
| C75% | Forelimb | 3.121806 | 0.001 | 0.67773 |
| plast | Forelimb | 1.692266 | 0.04 | 0.522806 |
| All | Forelimb (body mass adjusted) | 1.459471 | 0.08 | 0.679644 |
| C2 | Forelimb (body mass adjusted) | 0.644442 | 0.273 | 0.526224 |
| C25% | Forelimb (body mass adjusted) | 1.005085 | 0.149 | 0.58151 |
| C50% | Forelimb (body mass adjusted) | 1.067842 | 0.148 | 0.459029 |
| C75% | Forelimb (body mass adjusted) | 1.125707 | 0.144 | 0.441235 |
| plast | Forelimb (body mass adjusted) | 1.760136 | 0.037 | 0.542996 |
| All | Head mass | 3.643713 | 0.001 | 0.847379 |
| C2 | Head mass | 3.070694 | 0.002 | 0.712697 |
| C25% | Head mass | 3.028046 | 0.002 | 0.723063 |
| C50% | Head mass | 2.898176 | 0.001 | 0.608326 |
| C75% | Head mass | 2.73559 | 0.003 | 0.610777 |
| plast | Head mass | 2.694404 | 0.006 | 0.604032 |
| All | Head mass (body mass adjusted) | 1.371967 | 0.092 | 0.674756 |
| C2 | Head mass (body mass adjusted) | 1.801278 | 0.035 | 0.605788 |
| C25% | Head mass (body mass adjusted) | 2.052222 | 0.018 | 0.653079 |
| C50% | Head mass (body mass adjusted) | 0.478936 | 0.328 | 0.417745 |
| C75% | Head mass (body mass adjusted) | 1.76419 | 0.042 | 0.494712 |
| plast | Head mass (body mass adjusted) | 1.019314 | 0.161 | 0.47797 |
| All | Hindlimb | 3.779951 | 0.001 | 0.838382 |
| C2 | Hindlimb | 2.710314 | 0.002 | 0.685808 |
| C25% | Hindlimb | 3.101518 | 0.001 | 0.736733 |
| C50% | Hindlimb | 3.265116 | 0.001 | 0.644266 |
| C75% | Hindlimb | 2.664147 | 0.002 | 0.605541 |
| plast | Hindlimb | 1.842359 | 0.032 | 0.53528 |
| All | Hindlimb (body mass adjusted) | 1.004854 | 0.157 | 0.647228 |
| C2 | Hindlimb (body mass adjusted) | 0.926853 | 0.179 | 0.562345 |
| C25% | Hindlimb (body mass adjusted) | 0.682291 | 0.248 | 0.569105 |
| C50% | Hindlimb (body mass adjusted) | 0.878427 | 0.195 | 0.462769 |
| C75% | Hindlimb (body mass adjusted) | 0.251521 | 0.414 | 0.374447 |
| plast | Hindlimb (body mass adjusted) | 1.428475 | 0.085 | 0.521245 |
